# Biosynthesis of an Anti-Addiction Agent from the Iboga Plant

**DOI:** 10.1101/647891

**Authors:** Scott C. Farrow, Mohamed O. Kamileen, Lorenzo Caputi, Kate Bussey, Julia E. A. Mundy, Rory C. McAtee, Corey R. J. Stephenson, Sarah E. O’Connor

## Abstract

(−)-Ibogaine and (−)-voacangine are plant derived psychoactives that show promise as effective treatments for opioid addiction. However, these compounds are produced by hard to source plants making these chemicals difficult for broad-scale use. Here we report the complete biosynthesis of (−)-voacangine, and de-esterified voacangine, which is converted to (−)-ibogaine by heating. This discovery will enable production of these compounds by synthetic biology methods. Notably, (−)-ibogaine and (−)-voacangine are of the opposite enantiomeric configuration compared to the other major alkaloids found in this natural product class. Discovery of these biosynthetic enzymes therefore demonstrates how nature generates both enantiomeric series of this medically important alkaloid scaffold using closely related enzymes, including those that catalyze enantioselective formal Diels-Alder reactions.

**One Sentence Summary:** Biosynthesis of iboga alkaloids with anti-addiction promise reveals enantioselectivity of enzymatic Diels-Alder reactions.

## Main Text

Treatment of opiate addiction remains challenging, with over 45-thousand people in the United States dying in 2017 and a 500% increase in yearly opioid overdose deaths since the year 2000 (*1*). (−)-Ibogaine (**1**) (Fig. 1A), a plant-derived iboga-type alkaloid, has anti-addictive properties that were discovered accidentally by Howard Lotsof in 1962 when he noticed that ingesting this compound mitigated heroin cravings and acute opiate withdrawal symptomatology (*2*, *3*). Although the toxicity of (−)-ibogaine (**1**) has slowed its formal approval for addiction treatment in many countries, increased knowledge of its mode of action, side-effects and the discovery of (−)-ibogaine (**1**) analogs clearly indicate its potential as an anti-addictive agent (*2*–*4*). The plant that synthesizes (−)-ibogaine (**1**), *Tabernanthe iboga* (Iboga), is difficult to cultivate, prompting interest in developing biocatalytic methods for (−)-ibogaine (**1**) production. While the biosynthesis of the (+)-iboga type alkaloid scaffold has been recently elucidated (*5*), biosynthesis of the antipodal (−)-ibogaine (**1**) remained uncertain. Here we show that (−)-ibogaine (**1**) biosynthesis uses the same starting substrate as observed in (+)-iboga biosynthesis, but the key cyclization step proceeds *via* a distinct mechanism to generate the reduced iboga alkaloid (−)-coronaridine (**2**) (Fig. 1A). We further demonstrate that enzymatically generated (−)-coronaridine (**2**) can be 10-hydroxylated and 10-*O*-methylated (*6*) to form (−)-voacangine (**3**), and treatment of (−)-voacangine (**3**) with a *T. iboga* esterase reported here, followed by heating, yields (−)-ibogaine (**1**). This biocatalytic production strategy may facilitate sustainable and enhanced production of (−)-ibogaine (**1**) along with less toxic analogs, and moreover, reveals how two closely related enzyme systems generate two optical series *via* a formal Diels-Alder reaction.

**Fig 1.**
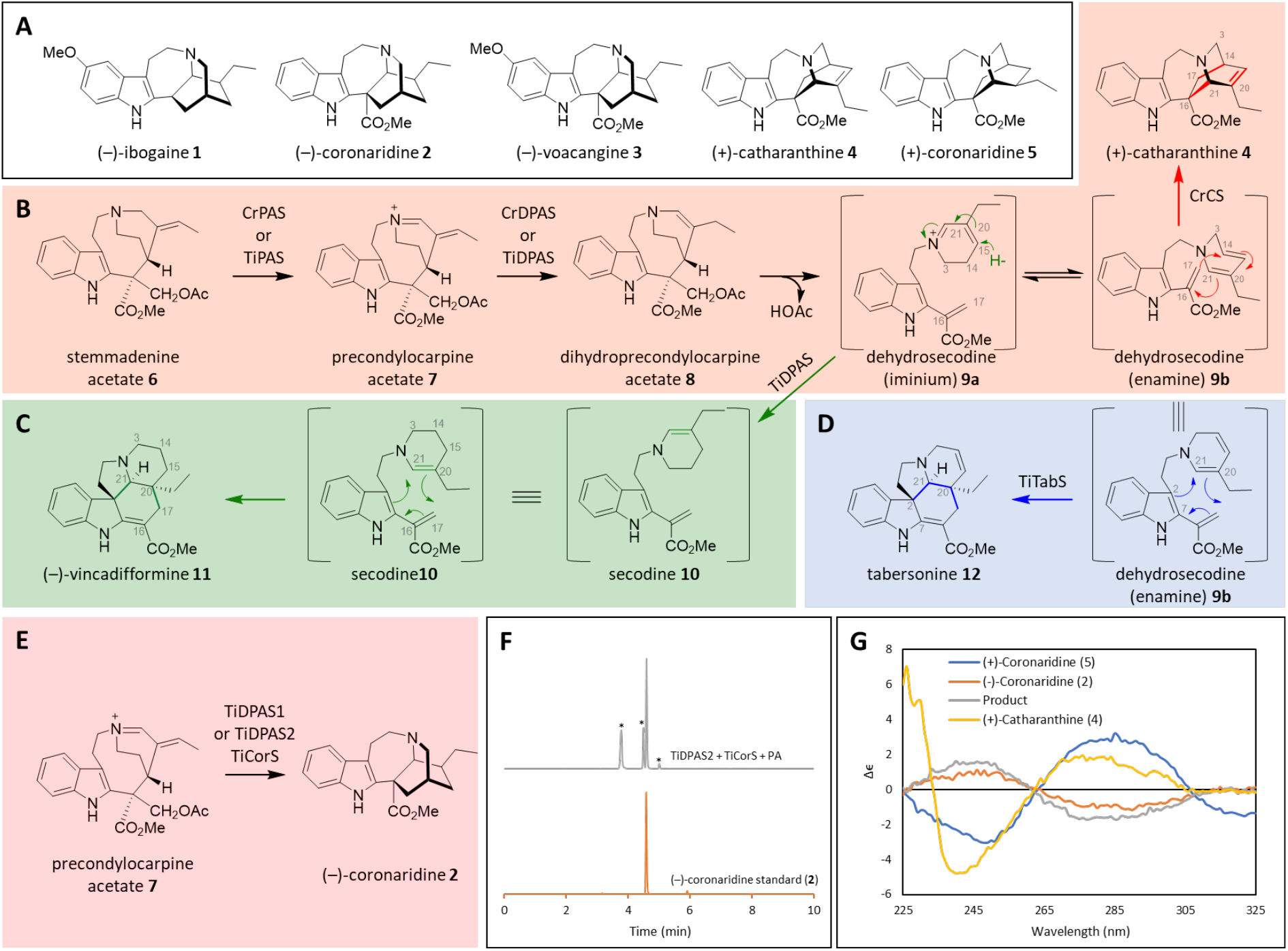
(+) and (−) iboga alkaloids. **A.** Anti-addiction agents (−)-ibogaine (**1**) and (−)-voacangine (**3**) are antipodal to (+)-catharanthine (**4**), a precursor to the anti-cancer drug vincristine. **B.** Biosynthesis of (+)-catharanthine (**4**). **C.** Biosynthesis of (−)-vincadifformine (**11**). **D**. Biosynthesis of (−)-tabersonine (**12**). **E.** Biosynthesis of the reduced iboga alkaloid (−)-coronaridine (**2**) directly from precondylocarpine acetate (**7**). **F.** LC-MS chromatogram showing formation of (−)-coronaridine (**2**) after incubation of precondylocarpine acetate (**7**) (50 μM) with TiDPAS2 (1 μM), TiCorS (5 μM) and NADPH (8 equivalents). Peaks marked with * are uncharacterized side products (*m/z* 339) that decomposed during isolation attempts. **G.** CD spectra of enzymatically produced coronaridine compared to authentic standards.

A transcriptome of *T. iboga* was previously obtained using seeds gifted by the Ibogaine Alliance (*6*). Although two late stage (−)-ibogaine (**1**) biosynthetic enzymes have been identified from this dataset, the biosynthetic pathway of the (−)-iboga scaffold remained unclear. Upon discovery of the pathway for the structurally related, antipodal iboga alkaloid (+)-catharanthine (**4**) from the plant *Catharanthus roseus* (Fig. 1A) (*5*), we hypothesized that *T. iboga* homologs of these *C. roseus* enzymes were responsible for biosynthesis of the (−)-iboga scaffold. (+)-Catharanthine (**4**) is synthesized by oxidation of stemmadenine acetate (**6**) by PAS (Precondylocarpine Acetate Synthase) to yield precondylocarpine acetate (**7**), which is reduced by DPAS (DihydroPrecondylocarpine Acetate Synthase) to give dihydroprecondylocarpine acetate (**8**). Dihydroprecondylocarpine acetate (**8**) desacetoxylates to form dehydrosecodine (**9**), which then undergoes a formal Diels-Alder cyclization catalyzed by CS (Catharanthine Synthase), an α/β hydrolase homolog, to yield (+)-catharanthine (**4**) (Fig. 1B, red box). We identified, cloned and heterologously expressed three homologs of PAS, two of DPAS and two of CS from the *T. iboga* transcriptome.

The three flavin-dependent PAS enzymes (TiPAS1, TiPAS2 and TiPAS3) each appeared to have the same biochemical function as the homolog involved in (+)-catharanthine (**4**) biosynthesis, CrPAS, namely oxidation of stemmadenine acetate (**6**) to precondylocarpine acetate (**7**) (Fig. 1B, Fig. S1). All subsequent experiments were performed with CrPAS, for which an optimized expression system had been developed (*5*). Next, TiDPAS1 (76.3% sequence identity to CrDPAS involved in catharanthine biosynthesis) and TiDPAS2 (86.3% sequence identity to CrDPAS) were tested with CrPAS and stemmadenine acetate (**6**). The expected reduced product, dihydroprecondylocarpine acetate (**8**) (Fig. 1B), is unstable and is not directly observed. In the presence of limiting amounts of the NADPH cofactor required by DPAS (1 substrate equivalent), a compound with *m/z* 337 was instead detected (Fig. S2). This product, which had been previously observed with CrDPAS (*5*), is likely an isomer of dehydrosecodine (e.g. **9a/9b**), which forms after desacetoxylation of dihydroprecondylocarpine acetate (**8**) (Fig. 1B). Surprisingly, in the presence of excess NADPH cofactor (>8 substrate equivalents), addition of either TiDPAS1 or TiDPAS2 generated the alkaloid vincadifformine (**11**) (Fig. S3–6), which is formed through Diels-Alder cyclization of reduced dehydrosecodine (secodine (**10**)) (Fig. 1C, green box). We hypothesize that with excess NADPH, TiDPAS1/2 over-reduces precondylocarpine acetate (**7**) to form secodine (**10**), which, in the absence of a dedicated cyclase enzyme, cyclizes spontaneously *via* a Diels-Alder mechanism to form vincadifformine (**11**). Biomimetic syntheses have shown that a (±)-vincadifformine analog readily forms from secodine (**10**), potentially due to the propensity of secodine (**10**) to adopt the conformation required for this Diels-Alder cyclization (*7*, *8*). The vincadifformine (**11**) observed in the CrPAS/TiDPAS reactions could be the result of a non-enzymatic cyclization, though surprisingly, CD spectra indicate that this reaction product is enantiomerically enriched (−)-vincadifformine (**11**) (Fig. S7). Alternatively, binding of secodine (**10**) to TiDPAS1/2 may provide an enantiomerically enriched cyclization product. However, formation of (−)-vincadifformine (**11**) is not dependent on the presence of a dedicated cyclase enzyme (*9*).

Finally, we tested the function of the two observed *T. iboga* CS homologs, which could be responsible for the key cyclization step to the (−)-iboga enantiomer. Addition of one of these homologs to assays containing stemmadenine acetate (**6**), CrPAS and TiDPAS1/TiDPAS2 and varying amounts of NADPH, yielded the alkaloid tabersonine (**12**), which forms through an alternative Diels-Alder cyclization mode of dehydrosecodine (**9**) (*5*) (Fig. 1D, blue box, Fig. S8), and this enzyme was thus named TiTabS (Tabernanthe iboga Tabersonine Synthase, 72.5 % sequence ID to previously identified tabersonine synthase from *C. roseus*, CrTS). *T. iboga* contains tabersonine (**12**) and numerous tabersonine (**12**) derived alkaloids (*10*), so identification of this enzyme activity, which has previously been observed in *C. roseus*, was expected. Incubation of stemmadenine acetate (**6**) with CrPAS, TiDPAS1/TiDPAS2 or precondylocaropine acetate (**7**) with TiDPAS1/TiDPAS2, excess NADPH and the second CS homolog from *T. iboga* led to the formation of the reduced iboga alkaloid coronaridine (**2**) (Fig. 1E), as evidenced by mass fragmentation and comparison to an authentic standard (Fig. 1F, Fig. S9–13). Although several side products were also observed in this *in vitro* enzymatic reaction (Fig. 1F), coronaridine (**2**) was the major product. This *T. iboga* CS homolog was named TiCorS (Coronaridine Synthase, 71.9 % sequence ID to CrCS). To assign the stereochemistry of coronaridine (**2**), the enzymatic product was isolated and subjected to CD analysis, which upon comparison to previous literature reports as well as authentic standards of (−)-coronaridine (**2**) (isolated from *Tabernaemontana divaricata*) (*6*) and (+)-coronaridine (**5**) (obtained from total synthesis, (*11*)) (Fig. 1G), indicated that the enzymatic product is (−)-coronaridine (**2**). Therefore, the biosynthetic pathways for both (+) and (−) iboga alkaloid scaffolds have now been elucidated.

We hypothesized how the (−)-coronardine enantiomer might form in this system. (−)-Catharanthine could be formed from dehydrosecodine (**9**), analogous to (+)-catharanthine (**4**) biosynthesis in *C. roseus* (*e.g.* Fig. 1B), and then subsequently reduced to form (−)-coronaridine (**2**). However, (−)-catharanthine is not observed as an intermediate in these enzymatic assays, nor has (−)-catharanthine been identified from natural sources. Alternatively, while it is obvious how vincadifformine (**11**) can be formed directly from secodine (**10**) (Fig. 1C), there is no logical mechanism by which secodine (**10**) can be cyclized to form (−)-coronaridine (**2**). Therefore we hypothesize that secodine (**10**) also does not act as an intermediate in (−)-coronaridine (**2**) biosynthesis. To gather evidence for an alternative mechanism, we isolated the product of precondylocarpine acetate (**7**) and TiDPAS1/2 that is formed under limiting NADPH conditions (**9**, *m/z* 337), which was also observed with DPAS from *C. roseus* (*5*). This compound **9** can be incubated with TiTabS to generate tabersonine (**12**), and thus can be inferred to be the dehydrosecodine substrate for the cyclases (Fig. S14). We incubated **9** with TiCorS, and, instead of observing the product directly as for tabersonine (**12**) biosynthesis, we observed an intermediate product, **13** (*m/z* 337). This product was isolated in partially purified form – though its instability prevented characterization – and then added to TiDPAS1/2 and NADPH (8 equivalents), resulting in the formation of (−)-coronaridine (**2**) (Fig. S15). Although both **9** and **13** were isolated in prohibitively low yield and were too unstable to be structurally characterized here, this experiment nevertheless suggests the appearance of distinct reaction intermediates. The reactivity of these intermediates suggests a reaction order and mechanism in which TiDPAS1/2 initially reduces precondylocarpine acetate (**7**) to generate dehydrosecodine (**9b**), which undergoes tautomerization (**9c-d,** (*12*, *13*)), followed by a formal [4+2] cyclization (*14*) to yield iminium **13**. Iminium **13** would then be reduced by TiDPAS1/2 to form (−)-coronaridine (**2**) (Fig. 2). In further support of this mechanism, when assays with **9**, TiCorS, TiDPAS2 and NADPH (8 equivalents) were performed in D2O, a compound co-eluting with (−)-coronaridine (**2**) with two additional atomic mass units was observed (Fig. S16). Notably, when CS and TS were assayed in D2O, no isotopic label was incorporated (Fig. S16). This provides further support that the mechanism of TiCorS, which requires tautomerization of the dihyropyridine ring, is distinct from the mechanism of CS and TS, which proceeds directly from dehydrosecodine (**9b**). This highlights that isomerization of dehydrosecodine provides the capacity for generating further structural diversity beyond (+)-catharanthine (**4**) and (−)-tabersonine (**12**). In short, this combination of cyclase and reductase generates an additional scaffold beyond the previously reported (+)-catharanthine (**4**) and (−)-tabersonine (**12**).

**Fig 2.**
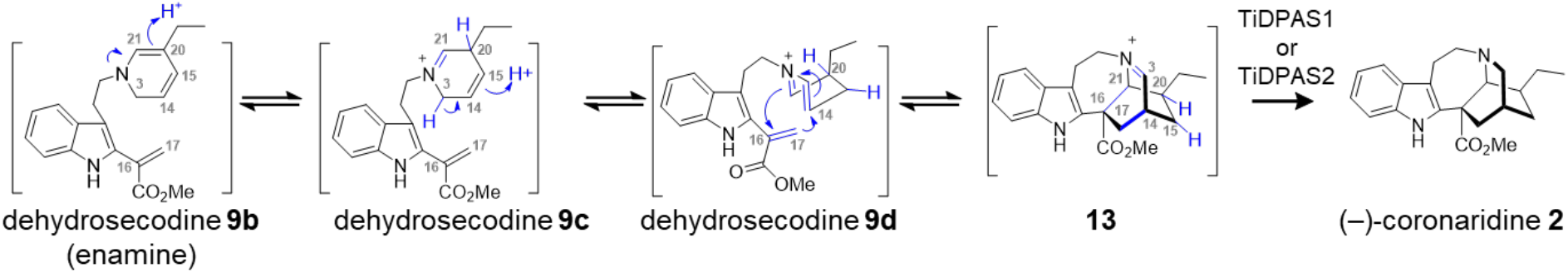
Biosynthesis of (−)-coronaridine (**2**) by TiDPAS1/2 and TiCorS. One plausible chemical mechanism for formation of the (−)-coronaridine (**2**) scaffold that is consistent with experimental evidence.

We next tested whether enzymatically synthesized (−)-coronaridine (**2**) could be used to generate (−)-voacangine (**3**) and (−)-ibogaine (**1**). We incubated the TiCorS/TiDPAS product with the previously identified ibogaine enzymes I10H and N10OMT, which respectively 10-hydroxylate and 10-*O*-methylate (−)-coronaridine (**2**) to generate (−)-voacangine (**3**)(*6*). Consistent with our structural assignment of (−)-coronaridine (**2**), incubation of the TiCorS/TiDPAS enzymatic product with these downstream enzymes yielded a product that was identical to an authentic standard of (−)-voacangine (**3**) (Fig. 3, Fig. S17). Not surprisingly, incubation of I10H and N10OMT with synthetic (+)-coronardine (**5**) gave only trace amounts of product (Fig. S18), highlighting the selectivity of the downstream enzymes to the stereochemistry of the substrate. In a well-established semi-synthetic process, (−)-voacangine (**3**) is subjected to a basic saponification to remove the methyl ester, and then heated to facilitate decarboxylation (*15*). The *T. iboga* transcriptome revealed three homologs of PNAE (Polyneuridine Aldehyde Esterase), an enzyme involved in the de-esterification and decarboxylation of ajmalan-type alkaloids found in other Apocynaceae plants (*16*), which had a similar co-expression profile as TiPAS1, TiDPAS2 and TiCorS. Incubation of TiPNAE1 with (−)-voacangine (**3**) yielded a product with a mass consistent with the de-esterified compound **15**, which then slowly converted to (−)-ibogaine (**1**) (Fig. 3). Heating of the de-esterified (−)-voacangine (**15**) product increased the efficiency and speed of this conversion (Fig. S19), suggesting that this decarboxylation is non-enzymatic, and a dedicated decarboxylase may be involved *in planta*. Regardless, enzymatic (−)-voacangine (**3**) provides an effective semi-synthetic starting material for (−)-ibogaine (**1**). Although TiPNAE1 could also de-esterify (−)-coronaridine (**2**), decarboxylation did not occur even after heating, suggesting that the electron donating methoxy group on the indole of (−)-voacangine (**3**) is required for spontaneous decarboxylation (Fig. S20).

**Fig 3.**
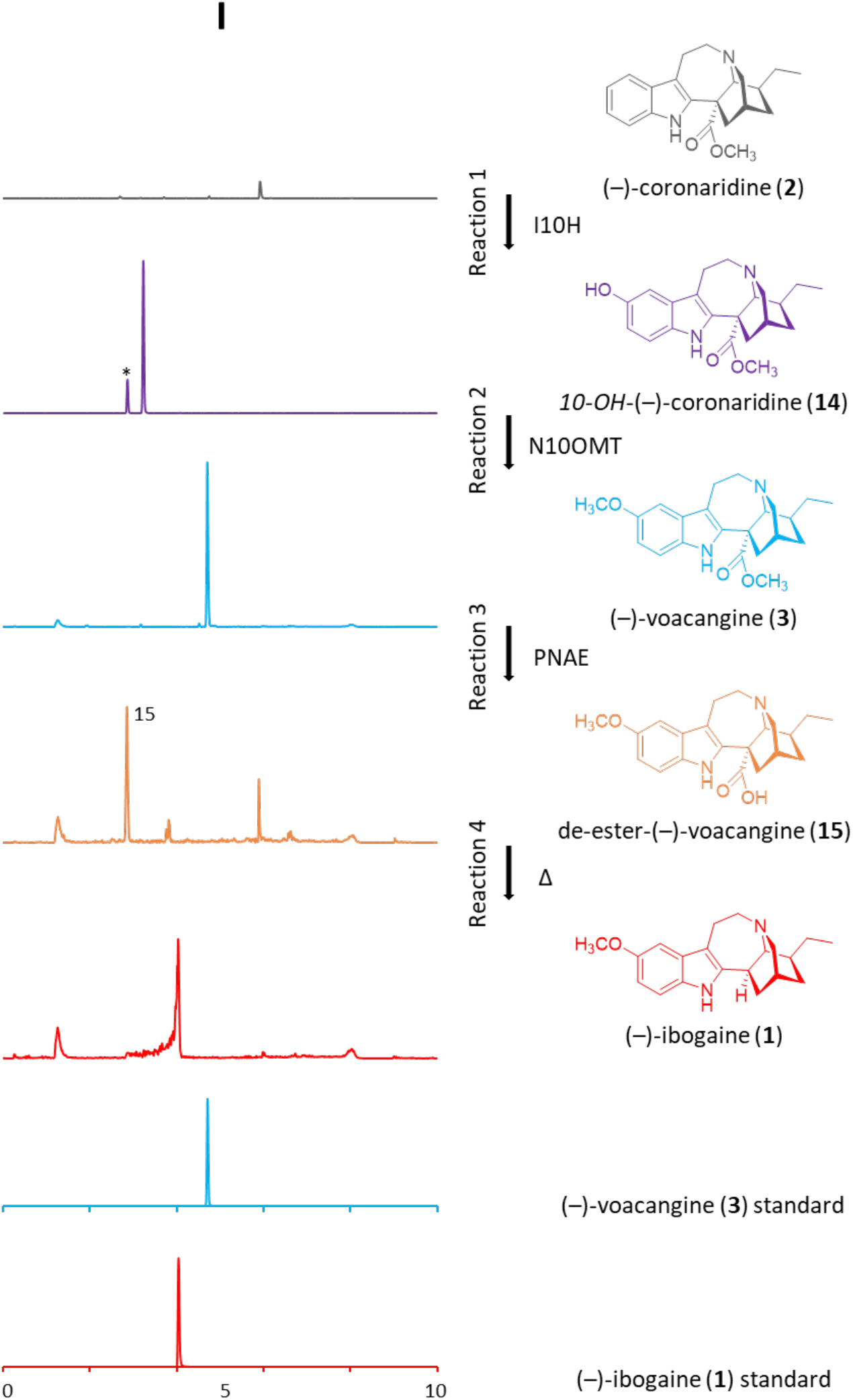
Formation of (−)-ibogaine (**1**) from enzymatically generated (−)-coronaridine (**2**). Peaks marked with * were products of endogenous yeast enzymes present in cultures expressing I10H.

(−)-Ibogaine (**1**) has a storied history dating back hundreds of years to the Congo Basin and the Bwiti religion, though it was the serendipitous discovery of the anti-addictive properties of (−)-ibogaine (**1**) that captured the attention of modern medicine. Here we report the biosynthesis of (−)-voacangine (**3**), and also report an esterase that may improve semi-synthesis of (−)-ibogaine (**1**). In addition to its extraordinary bioactivity, (−)-ibogaine (**1**) provides an opportunity to compare the formal Diels-Alder enzymatic synthesis of (−) and (+)-iboga enantiomers. The discovery and initial investigation of TiCorS suggest a possible mechanism for this cyclization and a basis for future enzyme engineering for this emerging class of cyclases. There are approximately 100 iboga alkaloids identified in nature (*10*), both of + and − optical series, and discovery of this pathway now provides full access to both medicinally important scaffolds (*17*), in addition to providing the first biocatalytic method for production of the anti-addictive alkaloid (−)-ibogaine (**1**).

## Acknowledgments

We gratefully acknowledge Dr. Kenneth Alper for introduction to this compound, and to Jonathan Dickerson of the Ibogaine Alliance for seeds. We thank Jakob Franke, Benjamin Lichman and Matthew DeMars for intellectual contributions, and Gerhard Saalbach and Carlo de-Oliveira-Martins for protein mass spectrometry. We thank David M. Lawson and Clare E. M. Stevenson for many helpful discussions.

## Author contributions

S.C.F. and S.E.O were responsible for conceptualization, methodology and writing; S.C.F., M.O.K., L.C., J.E.A.M., R.C.M., C.R.J.S. were responsible for analysis and investigation. J.E.A.M., S.C.F., M.O.K. performed CD spectroscopy, R.C.M., C.R.J.S. synthesized compounds.

## Data and materials availability

All data is available in the main text or the supplementary materials. Genbank accession numbers: MK840850 (TiPAS1); MK840851 (TiPAS2); MK840852 (TiPAS3); MK840853 (TiTabS); MK840854 (TiCorS); MK840855 (TiDPAS1); MK840856 (TiDPAS2); MK840857 (TiPNAE1); MK840858 (TiPNAE2); MK840859 (TiPNAE3).

## Supplementary Materials

### Materials and Methods

#### Chemicals and molecular biology reagents

All solvents used for extractions, chemical synthesis and preparative HPLC were HPLC grade, while solvents for UPLC/MS were MS grade. All solvents were purchased from Fisher Scientific. (−)-Ibogamine (**17**), (−)-ibogaine (**1**) and (−)-voacangine (**3**) were gifts from Dr. Kenneth Alper (New York University, USA). Authentic (−)-coronaridine (**2**) was extracted and purified from *Tabernaemontana divaricata* as previously described (*1*). (+)-Coronaridine (**5**) was synthesized by Professor Corey Stephenson and Rory McAtee (University of Michigan, USA). (+)-Catharanthine (**4**) was purchased from Sigma Aldrich. Tabersonine (**12**) was obtained from Ava Chem Scientific. Authentic (−)-vincadifformine (**11**) was provided by Dr. Vincent Courdavault (University of Tours, France) and Dr. Rod Andrade (Temple University, USA). Stemmadenine was purified by Professor Ivo J. Curcino Vieira as previously described (*2*). Stemmadenine acetate (**6**) was synthesized from stemmadenine by chemical acetylation as previously described (*2*).

Precondylocarpine acetate (**7**) was prepared enzymatically using the following conditions: 250 μg of stemmadenine acetate (**6**), 5 μg of CrPAS (GenBank MH213134) and 50 μg FAD were combined in a total reaction volume of 500 μL Tris-HCl buffer (50 mM, pH 8.5) and incubated at 37 °C for 2 h with shaking (1000 rpm). Reactions were quenched by addition of 5-volumes 90:9:1 methanol (MeOH):H_2_O:Formic acid (FA), and produced precondylocarpine acetate (**7**) was purified as previously described (*2*). Compound **9** was prepared enzymatically using the following conditions: 250 μg precondylocarpine acetate (**7**), 15 μg CrDPAS (GenBank KU865331) and 1 mg NADPH were combined in a total reaction volume of 500 μL Tris-HCl buffer (50 mM, pH 8.5) and incubated at 37 °C for 1 h with shaking (1000 rpm). Assays were quenched by addition of 5-volumes 90:9:1 MeOH:H_2_O:FA. Compound **9** was purified using a Dionex ultimate 3000 HPLC system equipped with a YMC-Pack Pro C18 column (250 × 10 mm, 5 μm, 12 nm) and UV detector monitoring 330 nm and was eluted in a gradient of 0.1 % FA and acetonitrile (MeCN) and immediately dried under vacuum using a Büchi roto-vap. Due to the unstable nature of **9**, assays were conducted immediately after purification.

Kanamycin sulfate, carbenicillin and gentamycin were from Formedium, and rifampicin was purchased from Sigma. All gene amplifications were performed using Platinum Superfi DNA polymerase (Thermo Fisher) and colony PCR was performed using Phire II master mix (Thermo Fisher). PCR product purification was performed using the Macherey-Nagel PCR clean-up kit. Plasmid purification was performed using the Promega Wizard miniprep kit. cDNA was prepared using Superscript IV VILO master mix and Turbo DNAse (Thermo Fisher).

#### Expression and purification of proteins

##### TiCorS, TiDPAS1, TiDPAS2, and TiPNAE1 expression in E. coli

*T. iboga* cDNA was isolated as previously described (*1*). Full-length TiCorS, TiDPAS1, TiDPAS2, and TiPNAE sequences were amplified from *T. iboga* cDNA using the primers listed. PCR products were purified from an agarose gel and ligated into the BamHI and KpnI restriction sites of the pOPINF vector (*3*) using the In-Fusion kit (Clontech Takara). Constructs were transformed into chemically competent *E. coli* Stellar cells and recombinant colonies were selected on LB agar + carbenicillin (100 μg/mL). Positive colonies were identified by colony PCR using primers and grown overnight at 37 °C for plasmid isolation and sequence confirmation using Eurofins sequencing service.

Chemically competent SoluBL21 *E. coli* cells (Amsbio) were transformed with appropriate plasmids by heat shock at 42 °C for 1 min. Transformed cells were selected on LB agar + carbenicillin (100 μg/mL). Single colonies were used to inoculate starter cultures of 50 mL 2 × YT medium supplemented with carbenicillin (100 μg/mL). Cultures were grown overnight at 37 °C with shaking (200 rpm). Starter cultures were used to inoculate 1 L of 2 × YT medium containing carbenicillin (100 μg/mL), grown to OD600=0.6 and then transferred to 16 °C for 30 min before inducing protein expression by addition of IPTG (0.3 mM). Protein expression was carried out for 16 h (18 °C, 200 rpm) where after cells were harvested by centrifugation (4000 × *g*) and re-suspended in 50 mL Buffer A (50 mM Tris-HCl pH 8, 50 mM glycine, 500 mM NaCl, 5% glycerol, 20 mM imidazole) with EDTA-free protease inhibitors (Roche Diagnostics Ltd.) and 10 mg lysozyme. Cells were lysed at 4 °C using a cell disruptor (25 KPSI) and centrifuged (35,000 × *g*) to remove insoluble debris. The supernatant was filtered through a 0.45 μm glass syringe filter (Sartorious) and applied to a HisTrap HP 5 mL column (GE Healthcare) with the assistance of an ÄKTA pure FPLC (GE Healthcare). Lysate was loaded at a flow rate of 2 mL/min and washed with Buffer A before being eluted with Buffer B (50 mM Tris-HCl pH 8, 50 mM glycine, 500 mM NaCl, 5% glycerol, 500 mM imidazole). Eluted proteins were further purified on a Superdex Hiload 16/60 S200 gel filtration column (GE Healthcare) at a flow rate of 1 mL/min using Buffer C (20 mM HEPES pH 7.5, 150 mM NaCl).

##### TiN10OMT and TiI10H

Cloning and expression of N10OMT and I10H was carried out as previously described (*1*).

##### Recombinant TiPAS from N. benthamiana

TiPAS1-3 full-length sequences were cloned into a modified TRBO vector (*4*) in which the cloning cassette of the pOPINF vector was inserted in the NotI restriction site. The vector was further modified by removing the *N*-terminal HisTag using PacI and SapI. PCR products were ligated into the modified vector upstream of the *C*-terminal HisTag using the In-fusion kit. Constructs were transformed into *E. coli* Stellar cells by heat shock (42 °C) and recombinant colonies were selected on LB agar + kanamycin (100 μg/mL). Positive colonies were screened by colony PCR and sequences were confirmed using Eurofins sequencing service. Sequence confirmed constructs were transformed into electrocompetent *Agrobacterium tumefaciens* strain GV3101 by electroporation. Recombinant colonies were selected on LB agar + rifampicin (100 μg/mL), gentamycin (50 μg/mL) and kanamycin (100 μg/mL). Single colonies were grown in 10 mL LB with antibiotics for 48 h at 28 °C and shaking (200 rpm). Cells were pelleted by centrifugation (4000 × *g*) and re-suspended in 10 mL infiltration buffer (10 mM NaCl, 1.75 mM CaCl2 and 100 μM acetosyringone). After incubation at room temperature for 3 h, cultures were diluted to OD600=0.1 and used to infiltrate 3-4-week old *N. benthamiana* plants. Leaves were harvested 5-days post-infiltration, and proteins were extracted from 30 g of tissue (fresh weight). Total proteins were extracted from pulverized leaf tissue with 100 mL ice-cold Tris-HCl buffer (50 mM, pH 8.0) containing EDTA-free protease inhibitors and 1 % PVPP. After incubation on ice for 1 h with intermittent vortexing, samples were filtered through miracloth and centrifuged (4000 × *g*) for 10 min to remove insoluble PVPP and tissue debris. The supernatant was further centrifuged (35,000 × *g*) for 20 min. Supernatants were collected and incubated with Ni-NTA agarose beads (Qiagen) equilibrated with Tris-HCl buffer (50 mM, pH 8.0) for 1 h. Samples were centrifuged (1,000 × *g*) for 1 min to pellet Ni-NTA agarose beads and washed 3 times with 10 mL Tris-HCl buffer (50 mM, pH 8). Proteins were eluted twice with 600 μL Tris-HCl (50mM, pH 8) containing 500 mM imidazole. Protein was buffer exchanged into HEPES (pH 7.5, 50 mM NaCl) and stored at –80 °C.

To confirm the presence of TiPAS1-3, proteins were subjected to trypsin digestion and LC/MS/MS analysis on a nano LC-orbitrap (see Data S1) with adaptations to (*5*). Protein samples were dissolved in 2.5 % sodium deoxycholate (Sigma, 30970-25G), pH 8.0, incubated with 10 mM DTT for 30 min at 65 °C followed by incubation with 20 mM iodoacetamide (IAA) at room temperature in the dark for 30 min, and quenched with 10 mM DTT (both in 50 mM TEAB). The samples were digested with sequencing grade trypsin (Promega) (1:50 w/w) and incubated at 50 °C for 8 h. Peptides were desalted using C18 tips, and aliquots were analyzed by nano LC-MS/MS on an Orbitrap Fusion Tribrid Mass Spectrometer coupled to an UltiMate 3000 RSLC nano LC system (Thermo Scientific, Hemel Hempstead, UK). The samples were loaded and trapped using a pre-column which was then switched in-line to the analytical column for separation. Peptides were separated on a nanoEase M/Z column (HSS C18 T3, 100 Å, 1.8 μm; Waters, Wilmslow, UK) using a gradient of MeCN at a flow rate of 0.25 μL/min with the following steps of solvents A (water, 0.1 % FA) and B (80 % MeCN, 0.1 % FA): 0-4 min 3 % B (trap only); 4-15 min increase B to 13 %; 15-77 min increase B to 38 %; 77-92 min increase B to 55 %; followed by a ramp to 99 % B and re-equilibration to 3 % B.

Data dependent analysis was performed using parallel CID and HCD fragmentation with the following parameters: positive ion mode, orbitrap MS resolution = 120k, mass range (quadrupole) = 300-1800 *m/z*, MS2 top20 in ion trap, threshold 1.9e4, isolation window 1.6 Da, charge states 2-5, AGC target 1.9e4, max inject time 35 ms, dynamic exclusion 1 count, 15 s exclusion, exclusion mass window ± 5 ppm. MS scans were saved in profile mode while MS2 scans were saved in centroid mode.

Recalibrated peak lists were generated with MaxQuant 1.6.3.4 (*6*) using *T. iboga* and *N. benthamiana* databases. The final database search was performed with the merged HCD and CID peak lists from MaxQuant using in-house Mascot Server 2.4.1 (Matrixscience, London, UK). The search was performed on the *T. iboga* and *N. benthamiana* protein sequences and the MaxQuant contaminants databases. For the search a precursor tolerance of 6 ppm and a fragment tolerance of 0.6 Da was used. The enzyme was set to trypsin/P with a maximum of 2 allowed missed cleavages. Oxidation (M) and deamidation (N/Q) were set as standard variable modifications and carbamido-methylation (CAM) of cysteine as fixed modification. The Mascot search results were imported into Scaffold 4.4.1.1 (www.proteomsoftware.com) using identification probabilities of 99 % for proteins and 95 % for peptides.

#### *In vitro* enzyme assays

##### TiPAS assays

*In vitro* assays with TiPAS1-3 were performed in 50 mM CHES buffer (pH 9.5), 50 μM FAD and 50 μM stemmadenine acetate (**6**). Due to low expression of recombinant TiPAS1-3, the amount of protein in these assays was not determined, however, 5 μL of HisTrap enriched protein was added to each assay. After TiPAS1-3 activity was demonstrated (Fig. S1), all subsequent assays were performed with purified CrPAS, for which an optimized expression system had already been established (*2*).

##### Coupled assays

Coupled assays were performed in 50 mM CHES buffer (pH 9.5), with CrPAS, 50 μM FAD and 50 μM stemmadenine acetate (**6**) (Fig. S2–4,8–10). In assays containing no CrPAS, and only TiDPAS1 or TiDPAS2 and TiCorS, 50 μM precondylocarpine acetate (**7**) was used in place of stemmadenine acetate (**6**) (Fig. S5–6,11–12). TiDPAS1 and TiDPAS2 require NADPH for activity. In assays producing putative dehydrosecodine isomer **9**, equimolar concentrations of NADPH were added (Fig. S2), whereas in assays producing (−)-vincadifformine (**11**) or (−)-coronaridine (**2**), a 1:8 ratio of substrate:NADPH was used (Fig. S3–6,9–12,15). The concentration of enzyme in coupled reactions was optimized as: 420 nM CrPAS; 1 μM TiPAS1 or TiDPAS2; 5 μM TiCorS or TiTabS. In assays commencing with TiCorS (Fig. S15A), 50 μM ca. of dehydrosecodine isomer **9** was added as substrate and incubated with 5 μM TiCorS for 20 min at 37 °C and shaking (1000 rpm), which yielded a product **13** with *m/z* 337 as determined using the UPLC/MS method described below (Fig S15A). Following analysis, **13** was isolated in partially purified form – though its instability prevented characterization– using the methods described for **9** and then added to TiDPAS1 or TiDPAS2 (1 μM) and NADPH (8 equivalents) for a further 20 min, resulting in the formation of (−)-coronaridine (**2**) (Fig. S15B). Assays were quenched by addition of 5-volumes 90:9:1 MeOH:H_2_O:FA and analyzed immediately using the UPLC/MS method described below.

For D_2_O experiments, assays were conducted in 50 mM CHES (pH 9.5) where ca. 50 μM dehydrosecodine (**9**) (that was isolated from reaction of precondylocarpine acetate (**7**) with DPAS), TiDPAS2 (1 μM) and TiCorS (5 μM) or TS or CS were added with excess NADPH (8-equivalents) to a total D_2_O or H_2_O reaction volume of 100 μL. Assays were incubated at 37 °C and shaking (1000 rpm) for 30 min and then quenched with 2-volumes 90:9:1 MeOH:H_2_O:FA. Assays were analyzed using the HR-MS method described below.

#### CD spectroscopy

Enzymatically prepared (−)-coronaridine (**2**) and (−)-vincadifformine (**11**) were generated using coupled assays in 50 mM CHES buffer (pH 9.5) for circular dichroism (CD) spectroscopy. A total of 10 mg stemmadenine acetate (**6**) was converted to (−)-coronaridine (**2**) or (−)-vincadifformine (**11**) using the coupled assay protocol described above, by pooling reactions in which 250 μg of stemmadenine acetate was used as starting material. Reactions were quenched by addition of 2 reaction volumes 90:9:1 MeOH:H_2_O:FA, and compounds were purified using an Agilent 1260 Infinity II HPLC system equipped with an Acquity BEH C18, 1.7 μm (2.1 × 50 mm) column. Separation was performed at 40 °C using a flow rate of 0.6 mL/min and 0.1 % ammonia as mobile phase A and MeCN as mobile phase B. Separation began with a 1 min isocratic step at 85 % A and 15% B followed by a linear gradient from 15 % to 60 % B in 2 min and 60 to 85 % B in 7.5 min. Conditions were then changed immediately to 100 % B for a 1 min wash followed by a return to initial conditions for a 1 min re-equilibration period prior to the next injection. Elution of compounds was monitored at 284 and 330 nm using an Agilent 1290 Infinity II PDA detector and fractions were collected using the fraction collector. Relevant fractions were pooled and dried under vacuum and stored at –80°C until analysis. Due to the unstable nature of intermediates under in-vitro assay conditions, recoveries of only ca. 50 μg each of **2** and **11** was achieved.

Products and standards were prepared to a final concentration of ≈200 μM in MS grade MeOH. Spectra were recorded in 1 nm steps with a 0.5 s averaging time on a Chirascan Plus spectropolarimeter (Applied Photophysics) at 20°C in a 0.5 mm cuvette. Measurements were collected in triplicates, averaged and background subtracted with MeOH.

#### Enzymatic synthesis of (−)-voacangine and semi-synthesis of (−)-ibogaine

Cultures of *S. cerevisiae* (1 mL) expressing I10H or empty vector were grown overnight at 30 °C with shaking (1000 rpm) in SC-leu media containing 2 % glucose. A 100 μL aliquot of this overnight culture was added to 900 μL buffered SC-leu media (pH 7.5, 50 mM HEPES) containing 2 % galactose to induce protein expression. Enzymatically prepared (−)-coronaridine (**2**), authentic isolated (−)-coronaridine (**2**) or authentic synthetic (+)-coronaridine (**5**) was added to separate I10H or empty vector expressing cultures to a final concentration of 50 μM. Reactions proceeded for 24 h at 30 °C and shaking (1000 rpm), where after cells were pelleted by centrifugation (4000 × *g*) and the supernatant was collected in a 15 mL falcon tube. Cell pellets were rinsed with 500 μL MeOH, sonicated and centrifuged (4000 × *g*) again to pellet debris. Supernatants were pooled and diluted to 10 mL with ultrapure water. Samples were purified on OASIS HLB cartridges (60 mg, 3 cc) that were activated and equilibrated with 3 mL MeOH and ultrapure water, respectively. Samples were loaded by gravity and washed with 3 mL ultrapure water followed by 3 mL 10 % MeOH. Compounds were eluted with 50:50 MeOH:MeCN v/v and dried under vacuum in a Genevac. Compounds were resuspended in 500 μL CHES buffer (50 mM, pH 9) and reacted with 100 ng of purified N10OMT and 1000 μM SAM for 2 h at 37 °C and shaking (1000 rpm). Purified TiPNAE1 (100 ng) was then added to these reactions and incubated under the same conditions for an additional 2 h. Reactions were quenched with 2-volumes MeOH and cleaned using the HLB protocol above. Eluents were dried under vacuum in a Genevac and analyzed using the UPLC/MS method (**A**) below.

#### Liquid-chromatography mass spectrometry analysis

##### UPLC/MS analysis of enzyme assays

Method **A**. For *in vitro* enzyme assays, compounds were separated on an Acquity BEH C18, 1.7 μm (2.1 × 50 mm) column (60 °C) using a Waters UPLC. Separation was performed using a flow rate of 0.6 mL/min and 0.1 % FA as mobile phase A and MeCN as mobile phase B. A linear gradient from 2 to 25 % B in 5 min and 25 to 62.5 % B in 7.5 min was applied for compound separation followed by an increase to 100 % B at 7.6 min. Conditions remained constant for 1 min and immediately changed to 2% B for a 1.3 min re-equilibration step prior to the next injection. Eluting compounds were subjected to positive ESI and analyzed on a Waters Xevo TSQ using optimized source conditions: Capillary voltage, 3.0 kV; source temperature, 150 °C; desolvation temperature, 450 °C; cone gas, 50 L/h; and desolvation gas, 800 L/h.

##### UPLC/MS assay of temperature-dependent (−)-voacangine decarboxylation

The effect of temperature on the decarboxylation of de-ester-(−)-voacangine (**15**) was carried out with modifications to UPLC/MS method (**A**) by increasing the column temperature in 5 °C increments at the beginning of each injection, up to 60 °C (Fig. S23).

##### HR-MS

For high resolution MS analysis (Fig. S16), compounds were separated on a Waters Acquity UPLC BEH C18 column (1 × 100 mm, 1.7 μm) in a gradient of 0.1% FA and MeCN and injected onto a Synapt G2 HDMS mass spectrometer (Waters) calibrated using a sodium formate solution. Samples were analyzed using positive ESI and monitored in the mass range of 50-1200 *m/z*. Capillary voltage was 3.5 V, cone voltage 40 V, source temperature 120 °C, desolvation temperature 350 °C, desolvation gas flow 800 L/h. Leu-enkephaline peptide (1 ng/μL) was used to generate a dual lock-mass calibration with [M+H]^+^ = 556.2766 and *m/z* =278.1135 measured every 10 sec. Spectra were generated in MassLynx 4.1 by combining a number of scans with background subtraction.

**Fig. S1.**
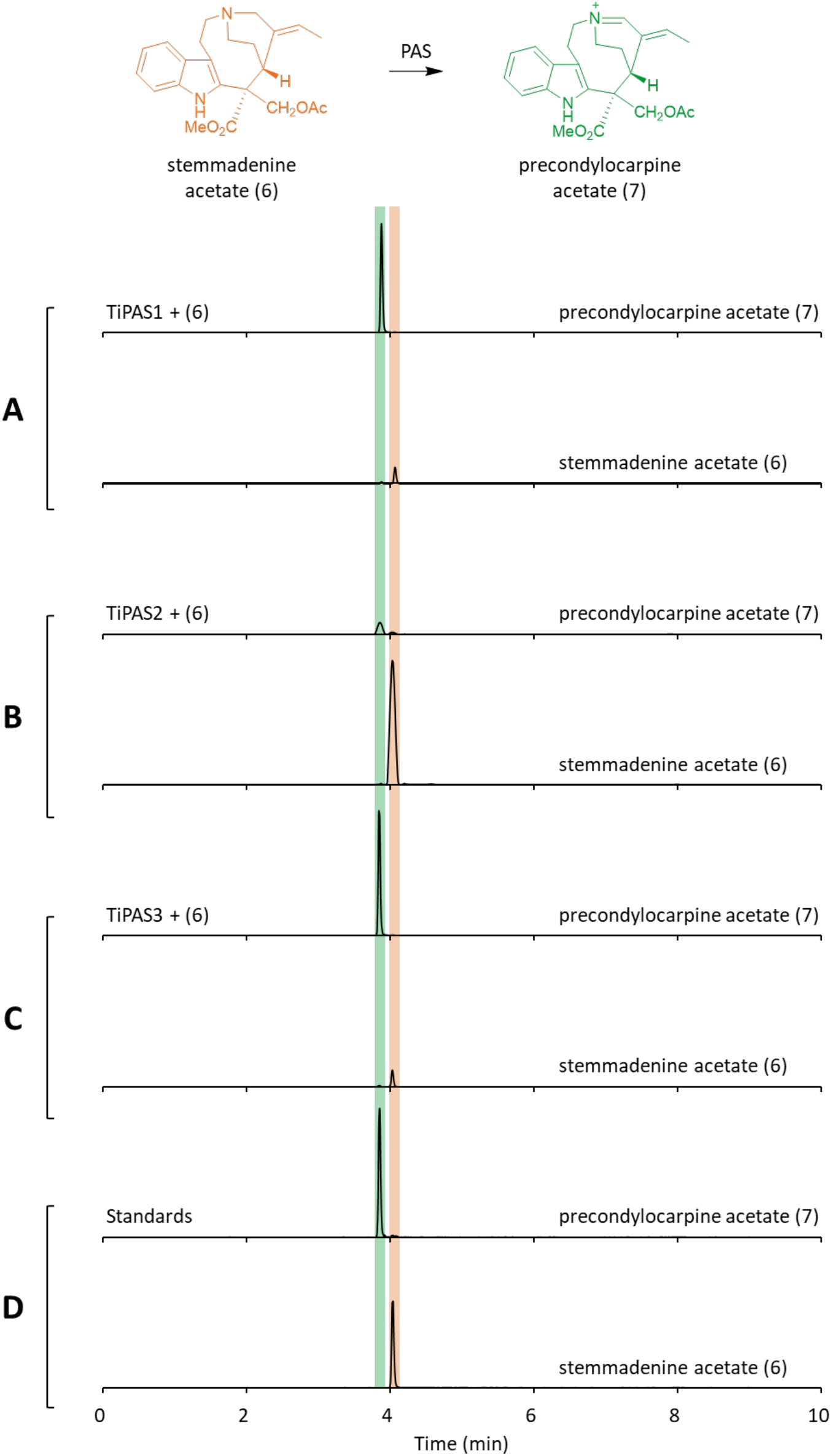
Formation of precondylocarpine acetate from stemmadenine acetate with TiPAS. **A.** UPLC/MS chromatograms illustrating the formation of precondylocarpine acetate (**7**, green) from stemmadenine acetate (**6**, orange) in assays with TiPAS1, **B.** TiPAS2, or **C.** TiPAS3. **D.** Authentic standards.

**Fig. S2.**
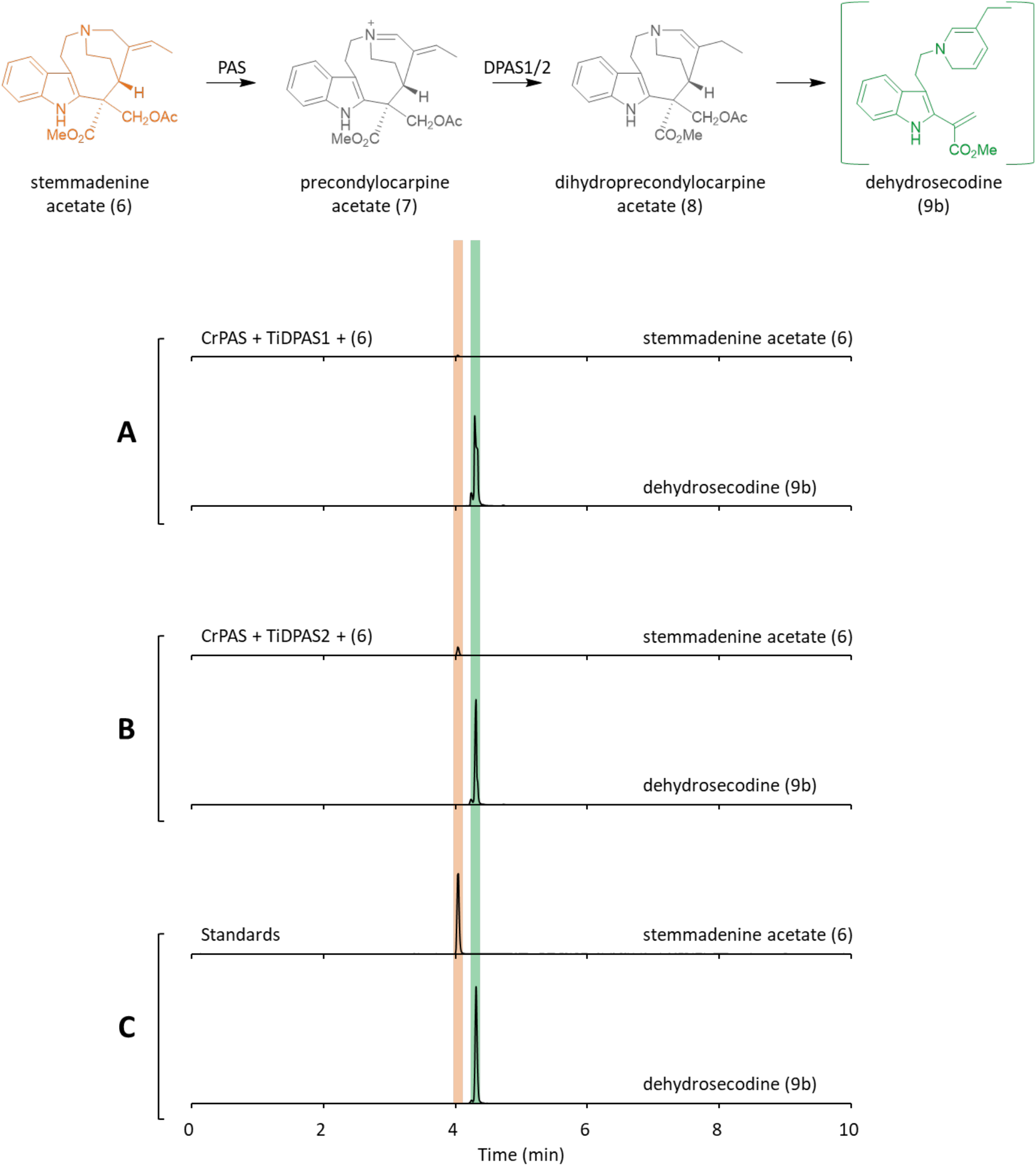
Formation of dehydrosecodine isomer from CrPAS, TiDPAS and stemmadenine acetate. **A.** UPLC/MS chromatograms illustrating the formation of putative dehydrosecodine isomer (**9**, green) from stemmadenine acetate (**6**, orange) in assays with CrPAS and TiDPAS1 or **B.** CrPAS and TiDPAS2. **C.** Authentic standards. Dehydrosecodine was not characterized due to low isolation yield and lability.

**Fig. S3.**
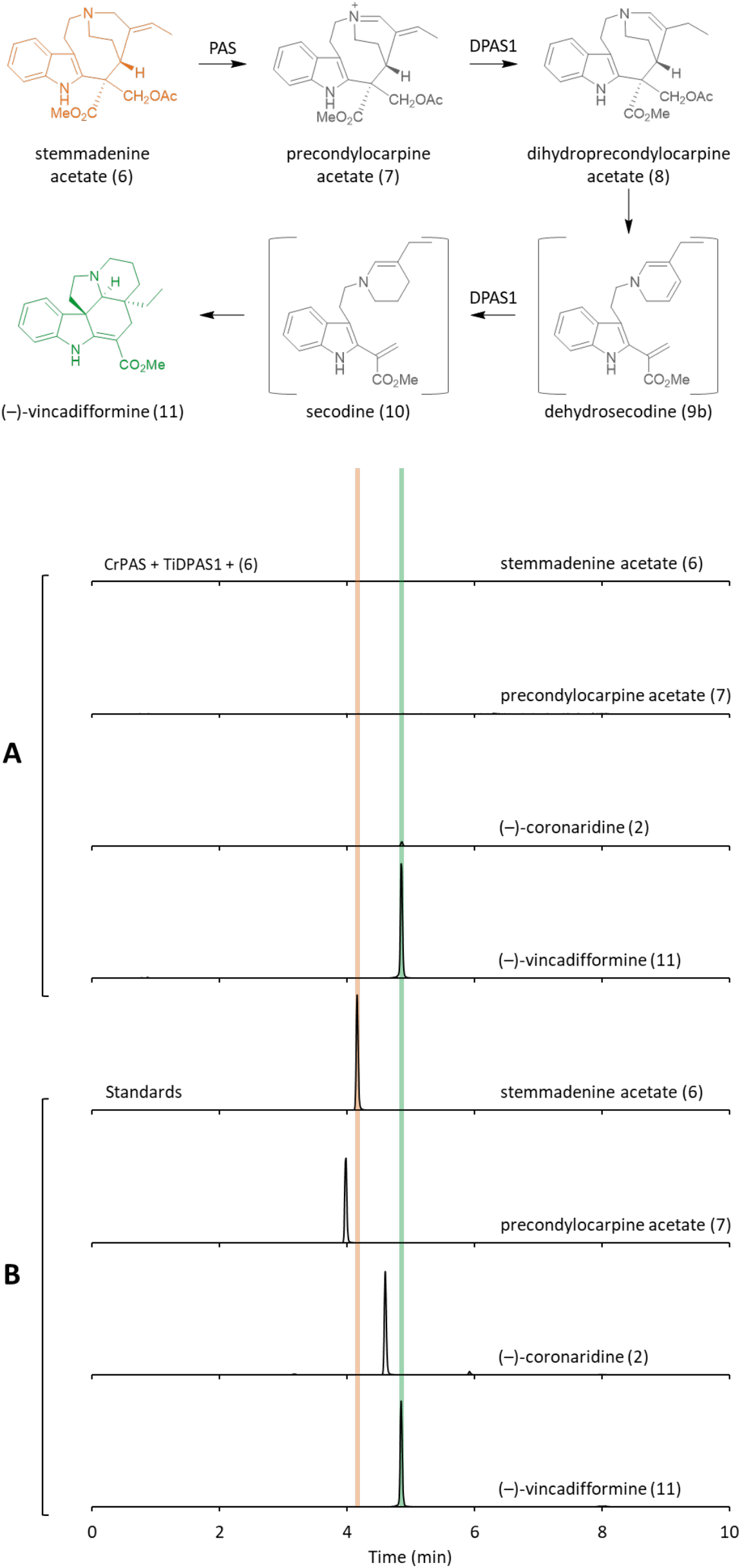
Formation of vincadifformine from stemmadenine acetate, CrPAS and TiDPAS1. **A.** UPLC/MS chromatograms illustrating the formation of (−)-vincadifformine (**11**, green) from stemmadenine acetate (**6**, orange) in assays with CrPAS and TiDPAS1. **B.** Authentic standards.

**Fig. S4.**
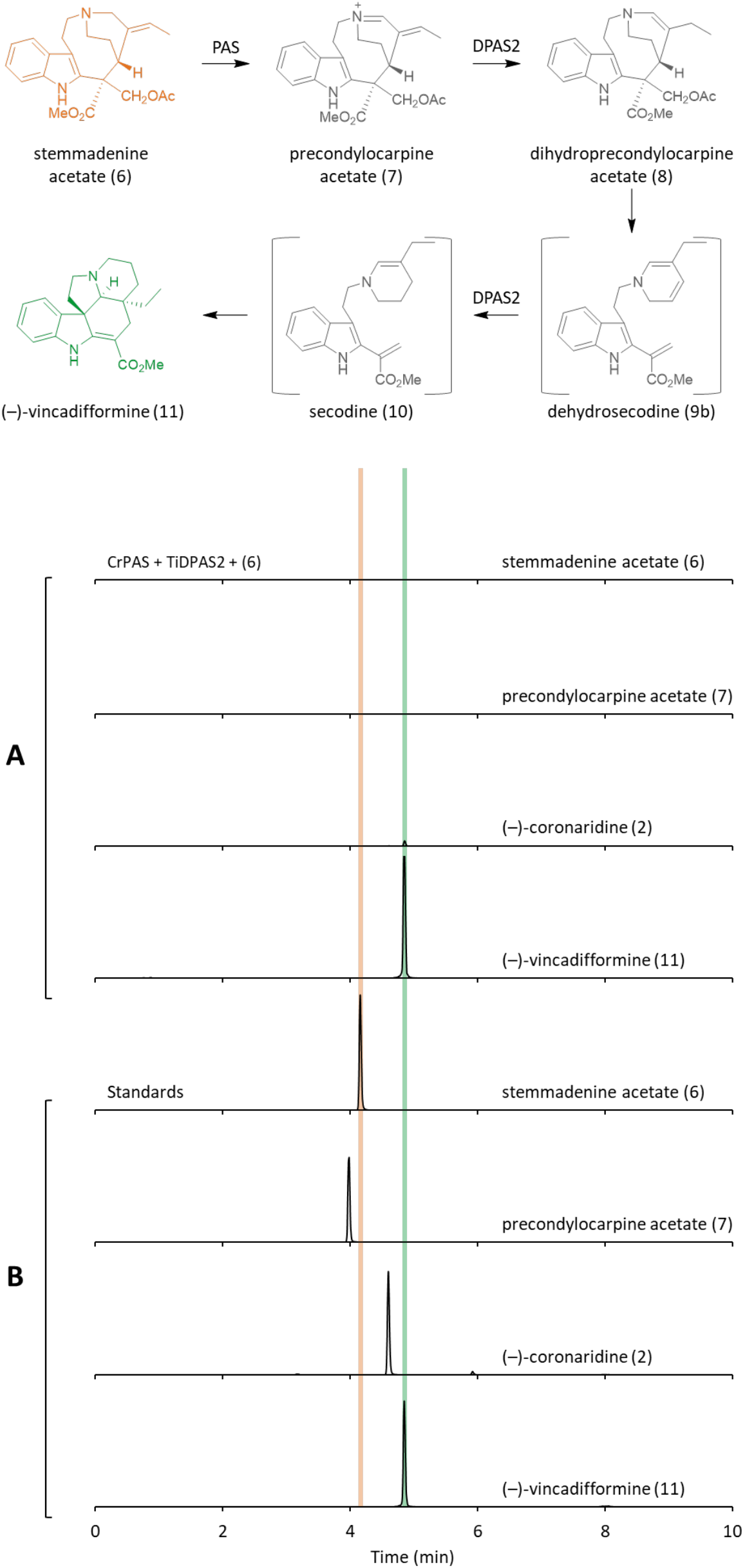
Formation of vincadifformine from stemmadenine acetate, CrPAS and TiDPAS2. **A.** UPLC/MS chromatograms illustrating the formation of vincadifformine (**11**, green) from stemmadenine acetate (**6**, orange) in assays with CrPAS and TiDPAS2. **B.** Authentic standards.

**Fig. S5.**
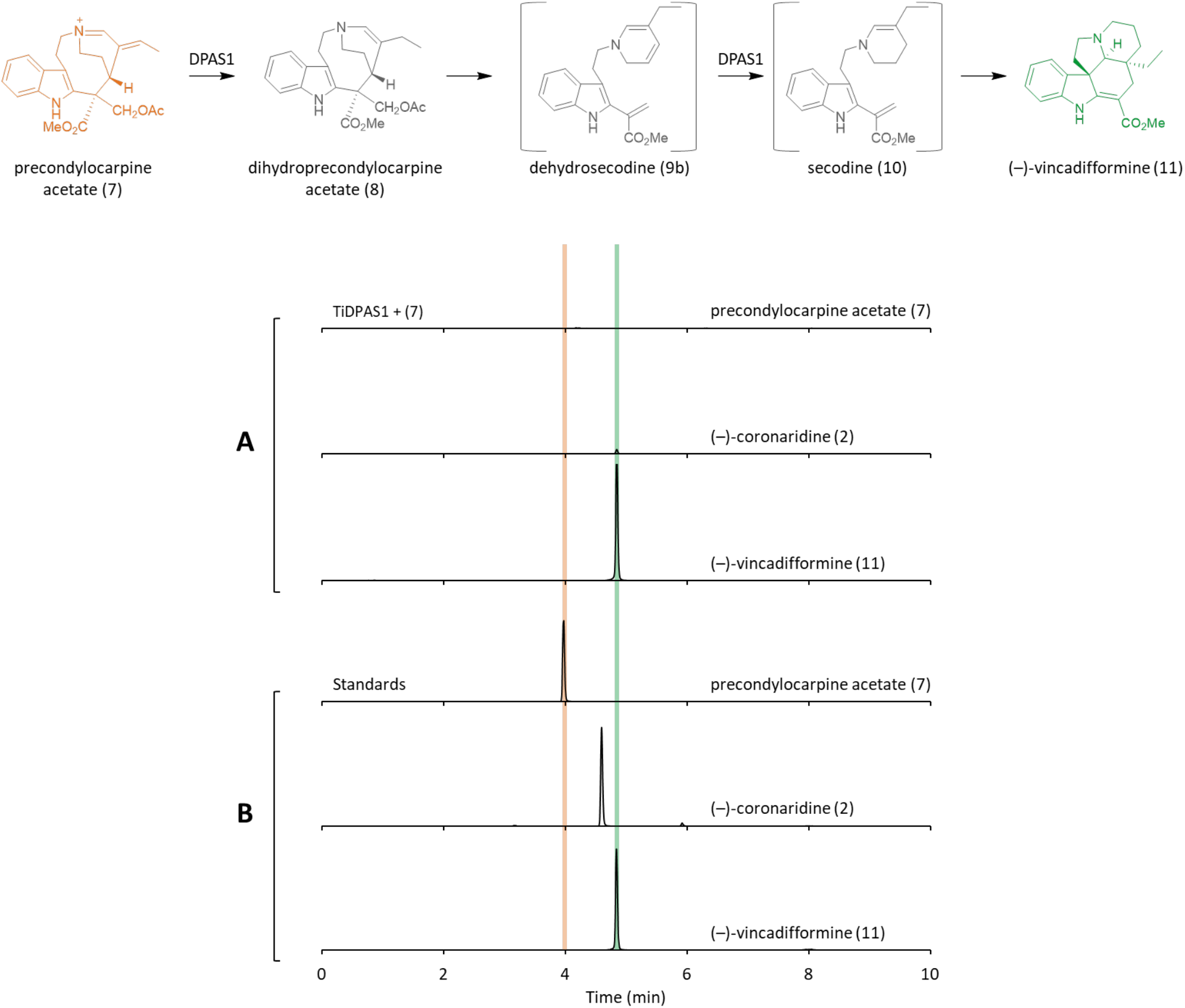
Formation of vincadifformine from precondylocarpine acetate and TiDPAS1. **A.** UPLC/MS chromatograms illustrating the formation of vincadifformine (**11**, green) from precondylocarpine acetate (**7**, orange) in assays with CrPAS and TiDPAS1. **B.** Authentic standards.

**Fig. S6.**
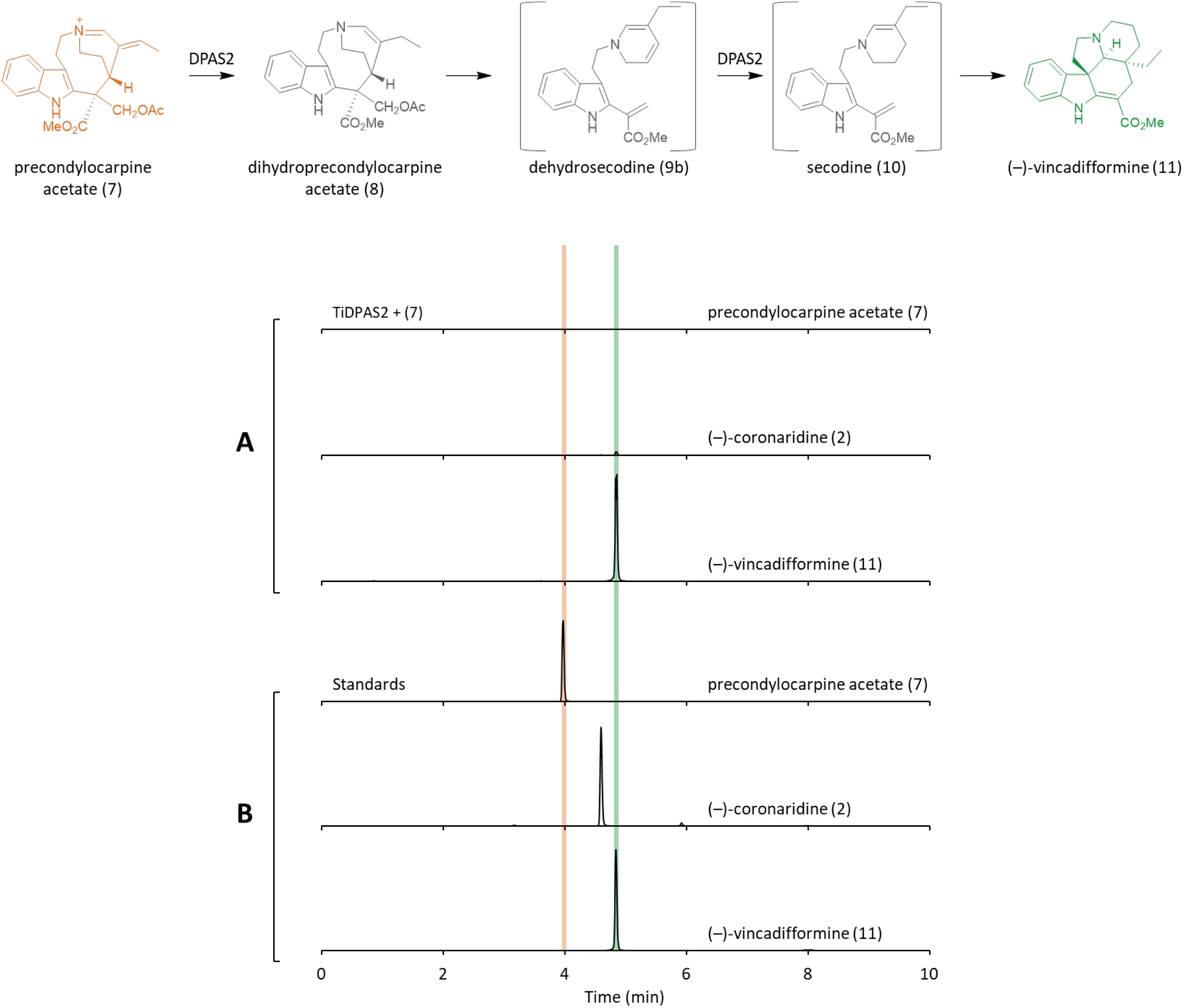
Formation of vincadifformine from precondylocarpine acetate and TiDPAS2. **A.** UPLC/MS chromatograms illustrating the formation of vincadifformine (**11**, green) from precondylocarpine acetate (**7**, orange) in assays with CrPAS and TiDPAS2. **B.** Authentic standards.

**Fig. S7.**
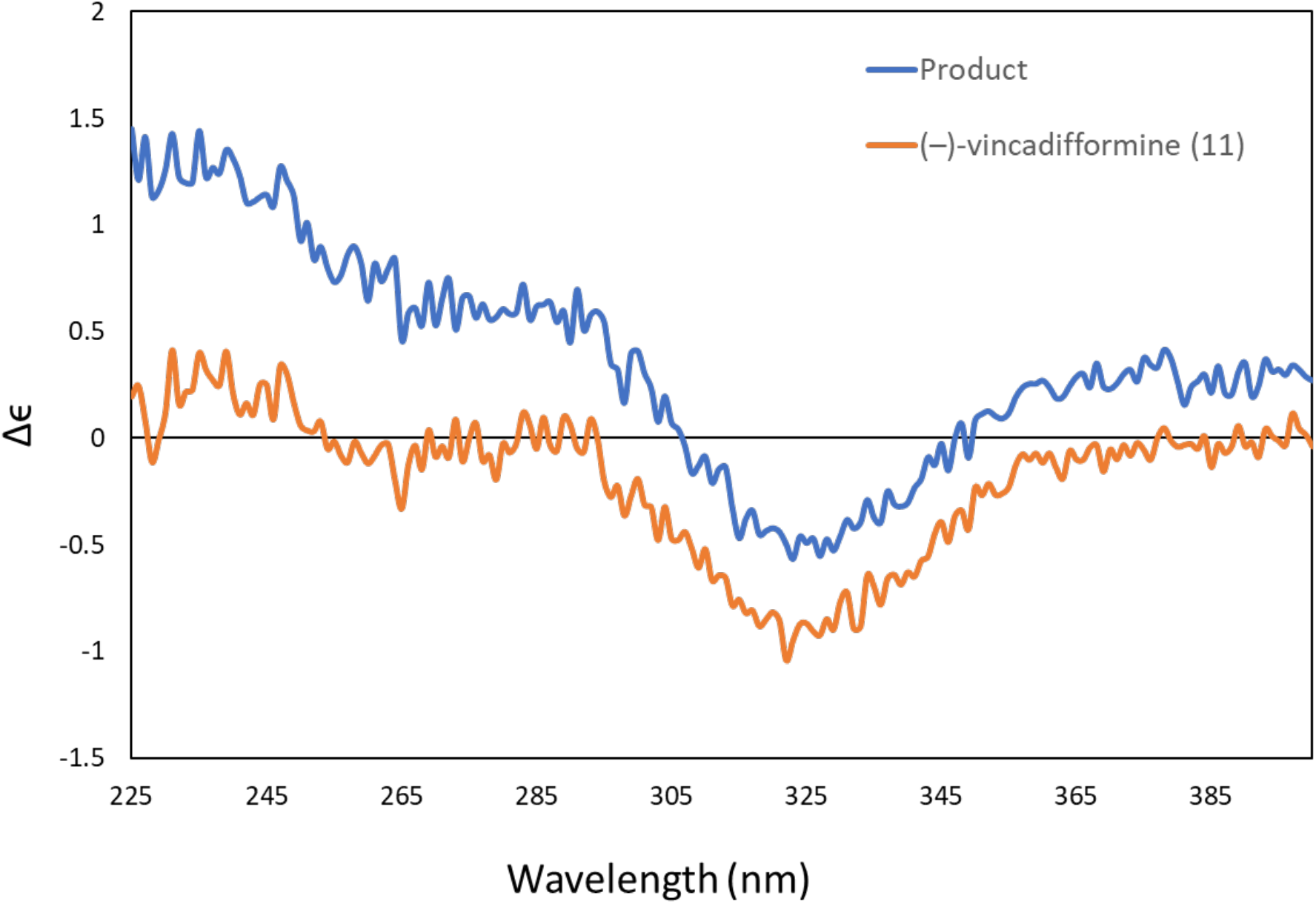
Stereochemistry of vincadifformine observed in enzyme assays. Stereochemical assignment of enzymatically produced vincadifformine (**11**, blue) using CD spectroscopy and comparison to an authentic (−)-vincadifformine standard (**11**, orange). Spectra were acquired in MS grade MeOH at a concentration of ≈200 μM.

**Fig. S8.**
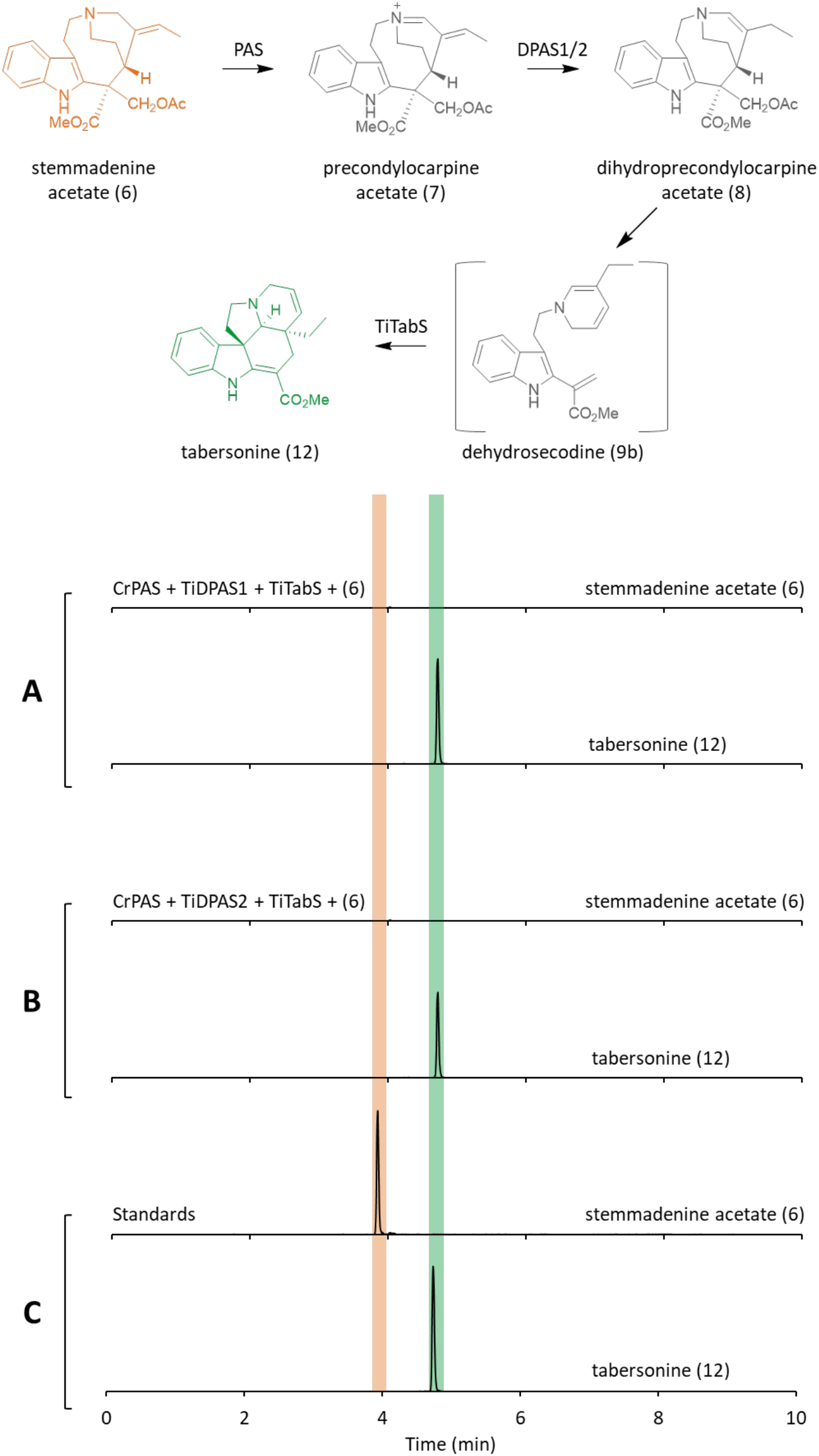
Formation of tabersonine from stemmadenine acetate. **A.** UPLC/MS chromatograms illustrating the formation of tabersonine (**12**, green) from stemmadenine acetate (**6**, orange) in assays with CrPAS, TiDPAS1 and TiTabS, or **B.** CrPAS, TiDPAS2 and TiTabS. **C.** Authentic standards.

**Fig. S9.**
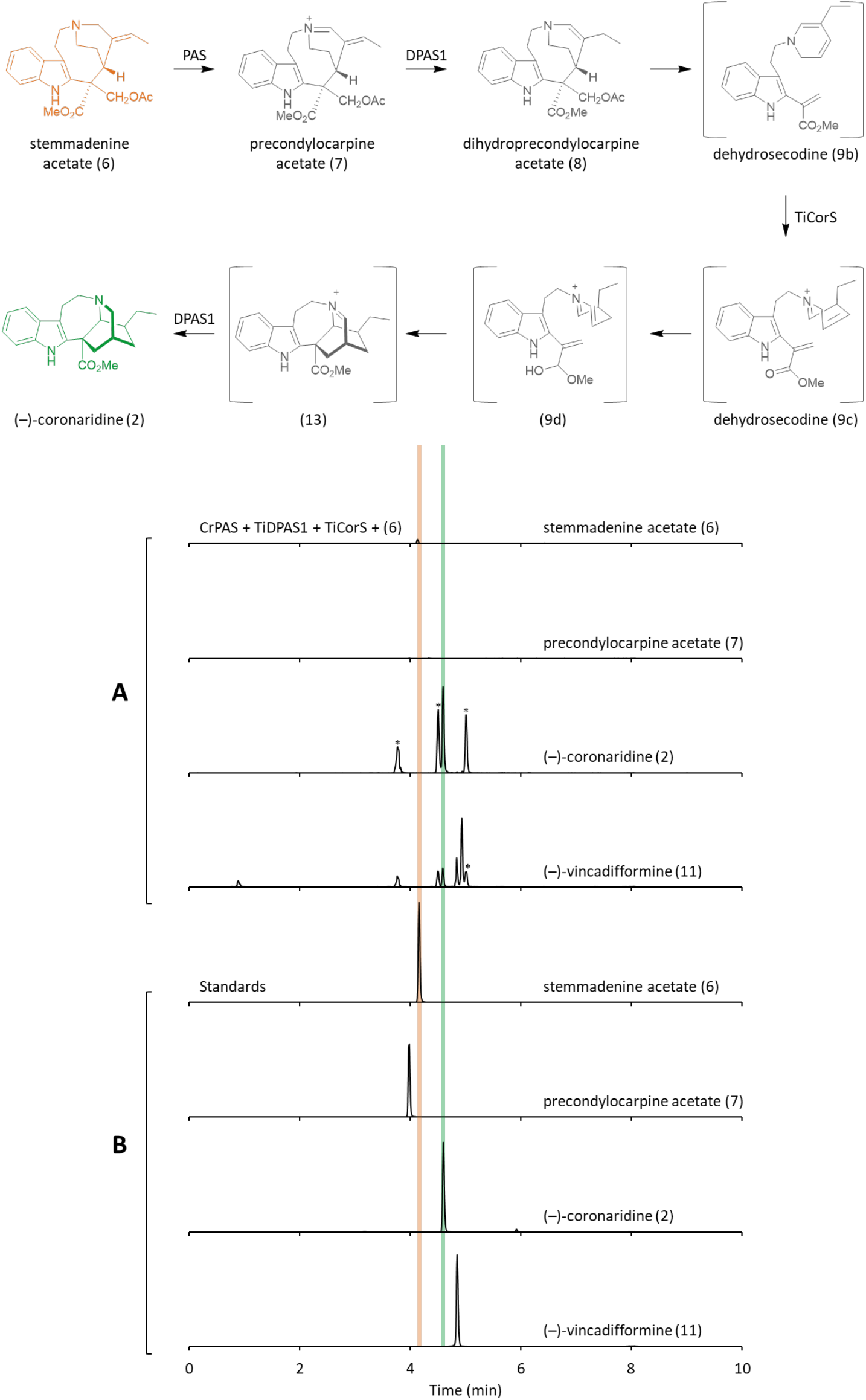
Formation of (−)-coronaridine from stemmadenine acetate (TiDPAS1). **A.** UPLC/MS chromatograms illustrating the formation coronaridine (**2**, green) from stemmadenine acetate (**6**, orange) in assays with CrPAS, TiDPAS1 and TiCorS. Peaks marked with * were labile, decomposed during isolation attempts and thus were not characterized. Trace quantities of vincadifformine (**11**) were detected. **B.** Authentic standards.

**Fig. S10.**
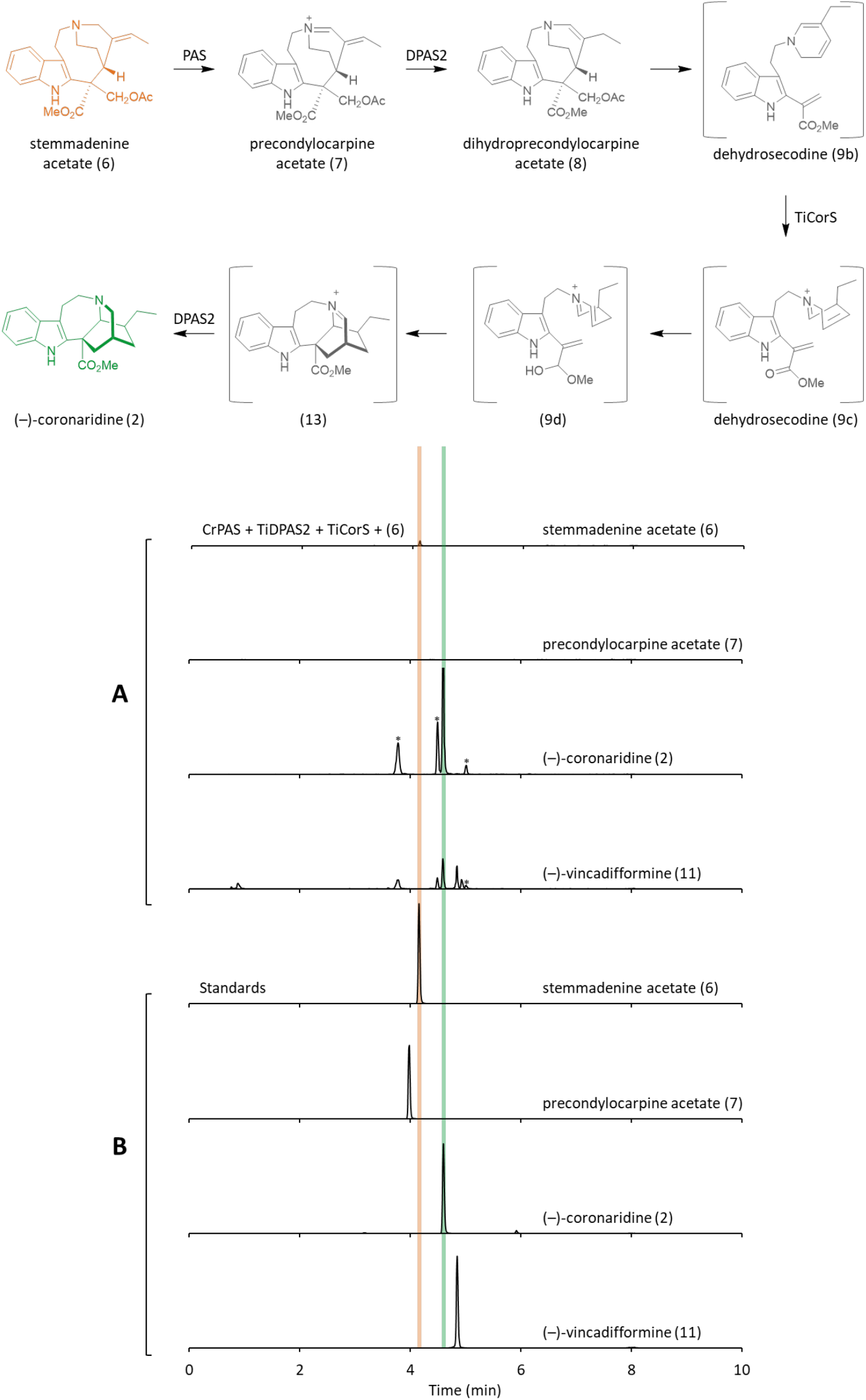
Formation of (−)-coronaridine from stemmadenine acetate (TiDPAS2). **A.** UPLC/MS chromatograms illustrating the formation coronaridine (**2**, green) from stemmadenine acetate (**6**, orange) in assays with CrPAS, TiDPAS2 and TiCorS. Peaks marked with * were labile and not characterized. Trace quantities of vincadifformine (**11**) were detected. **B.** Authentic standards.

**Fig. S11.**
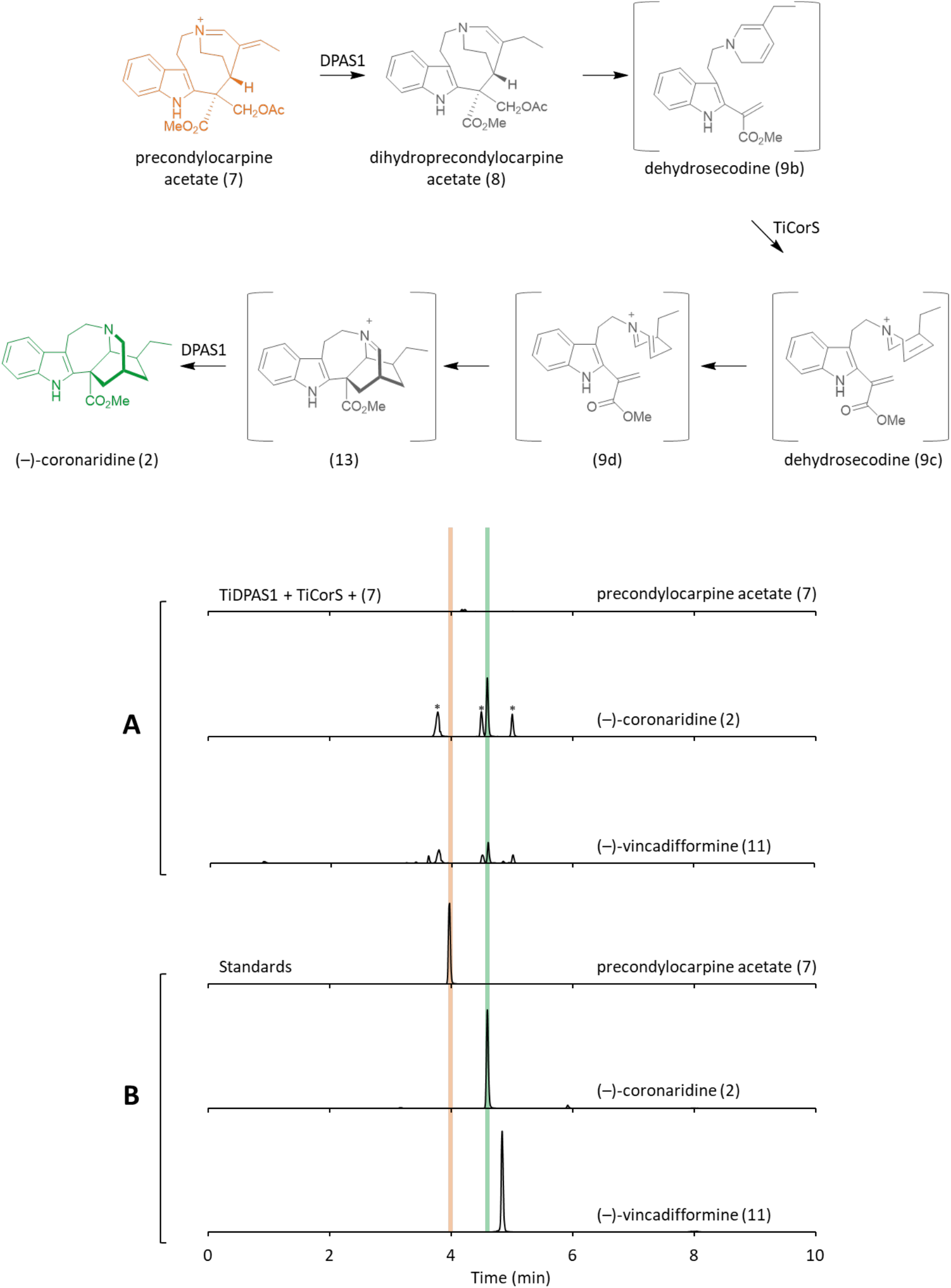
Formation of (−)-coronaridine from precondylocarpine acetate (TiDPAS1). **A.** UPLC/MS chromatograms illustrating the formation of coronaridine (**2**, green) from precondylocarpine acetate (**7**, orange) in assays with TiDPAS1 and TiCorS. Peaks marked with * were labile and not characterized. Trace quantities of vincadifformine (**11**) were detected. **B**. Authentic standards.

**Fig. S12.**
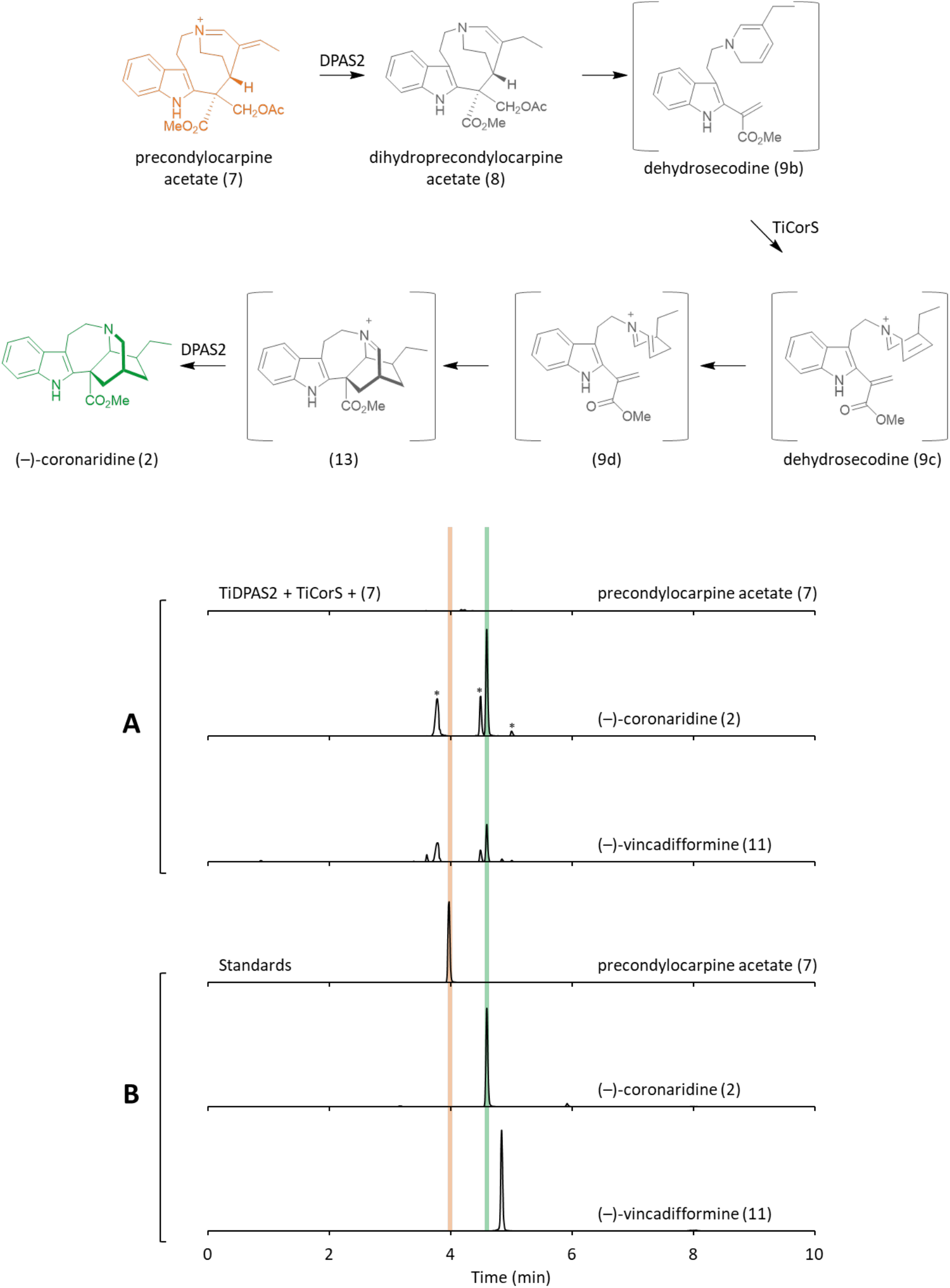
Formation of (−)-coronaridine from precondylocarpine acetate (TiDPAS2). **A.** UPLC/MS chromatograms illustrating the formation of coronaridine (**2**, green) from precondylocarpine acetate (**7**, orange) in assays with TiDPAS2 and TiCorS. Peaks marked with * were labile and not characterized. Trace quantities of vincadifformine (**11**) were detected. **B**. Authentic standards.

**Fig. S13.**
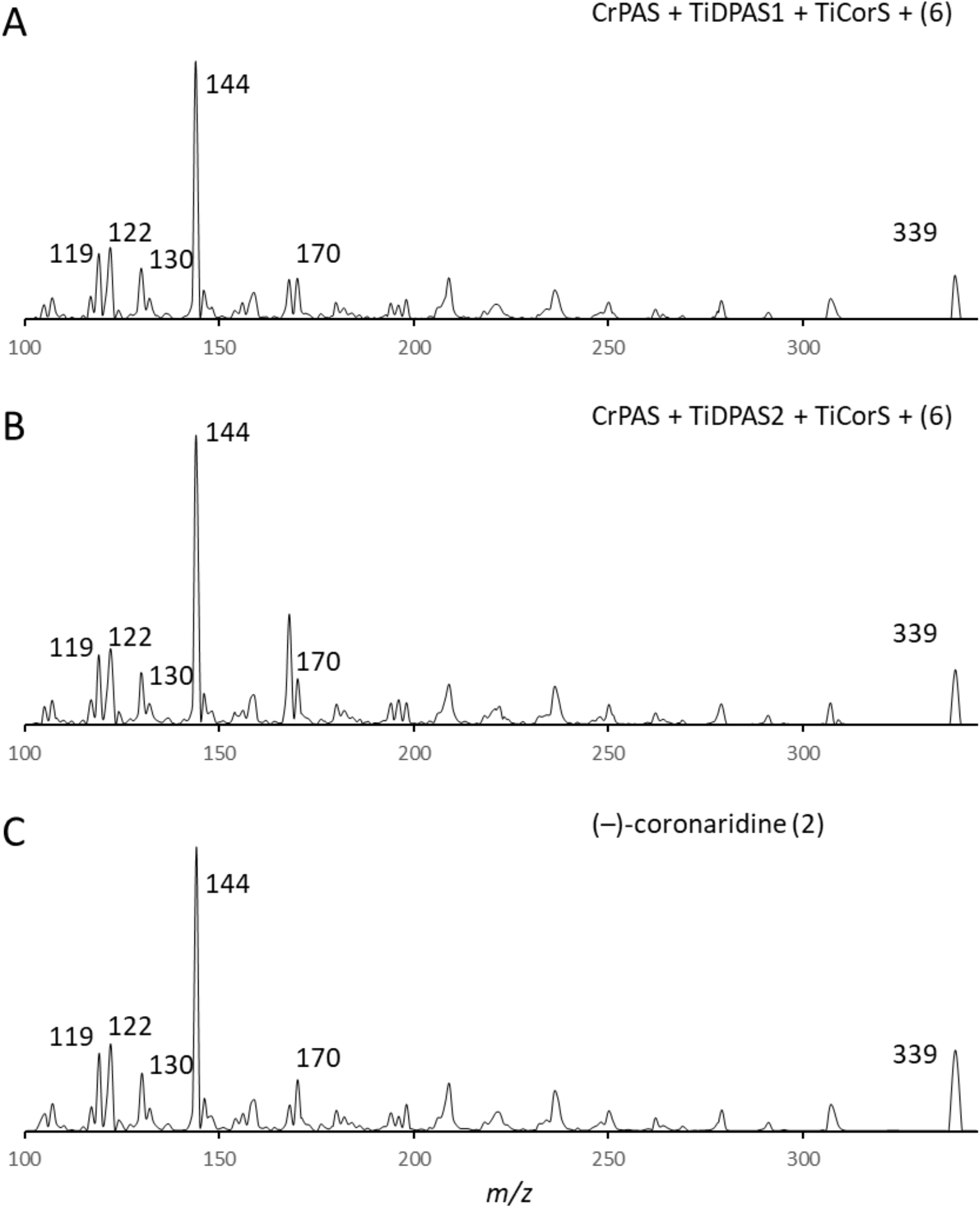
Product ion spectra of (−)-coronardine. **A.** MS/MS product ion spectra of enzymatically produced coronaridine (**2**) from assays containing CrPAS, TiDPAS1 and TiCorS, or **B.** CrPAS, TiDPAS2 and TiCorS. **C.** Product ion spectra of authentic (−)-coronaridine (**2**) standard.

**Fig. S14.**
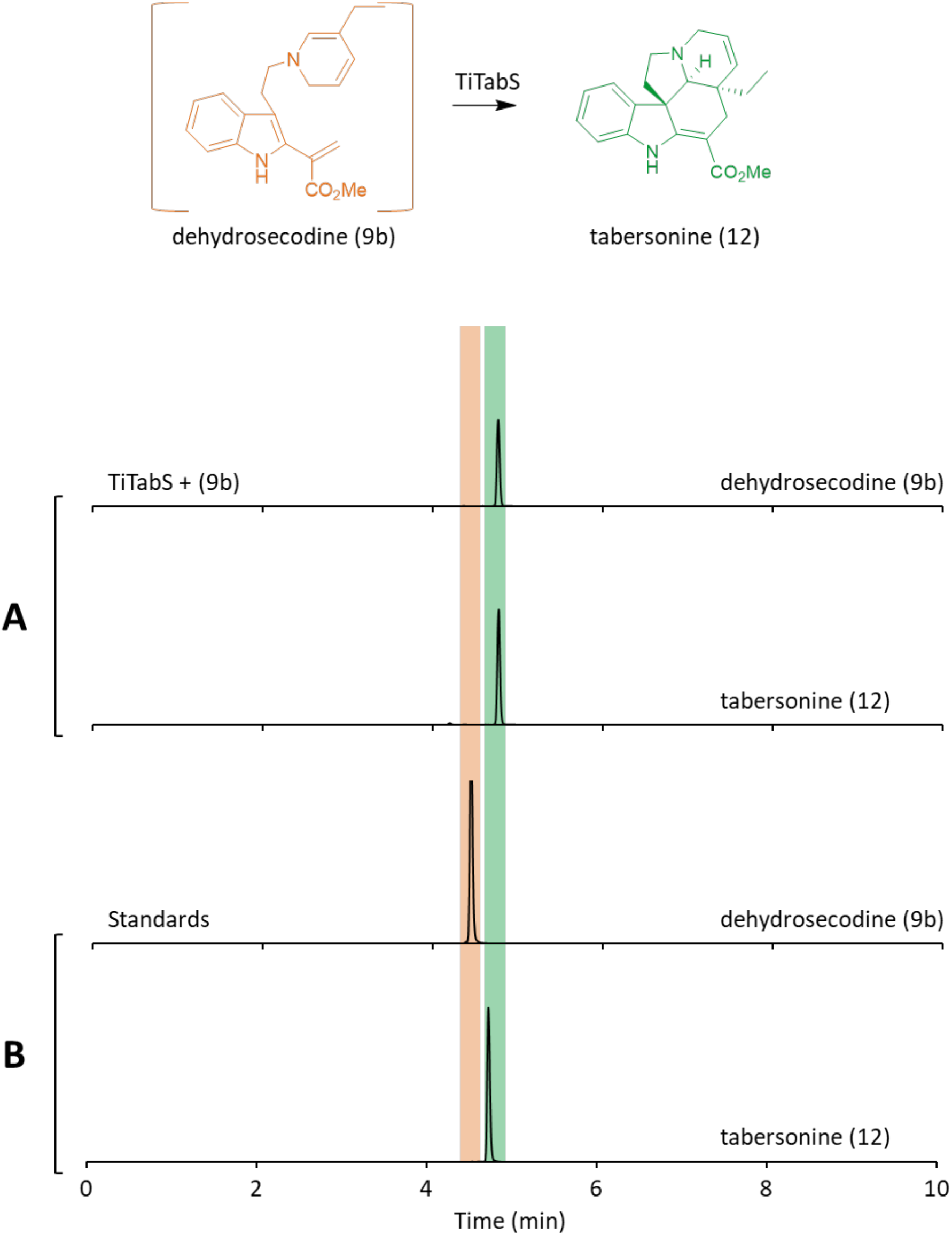
Activity of TiTabS from intermediate 9. **A.** UPLC/MS chromatograms illustrating the formation of tabersonine (**12**, green) from putative dehydrosecodine isomer **9**(orange), isolated from reaction of precondylocarpine acetate (**7**) and TiDPAS1) in assays with TiTabS. **B.** Authentic standards of **9**(isolated from reaction of **7**, TiDPAS1 and 1 eq. NADPH) and tabersonine (**12**).

**Fig. S15.**
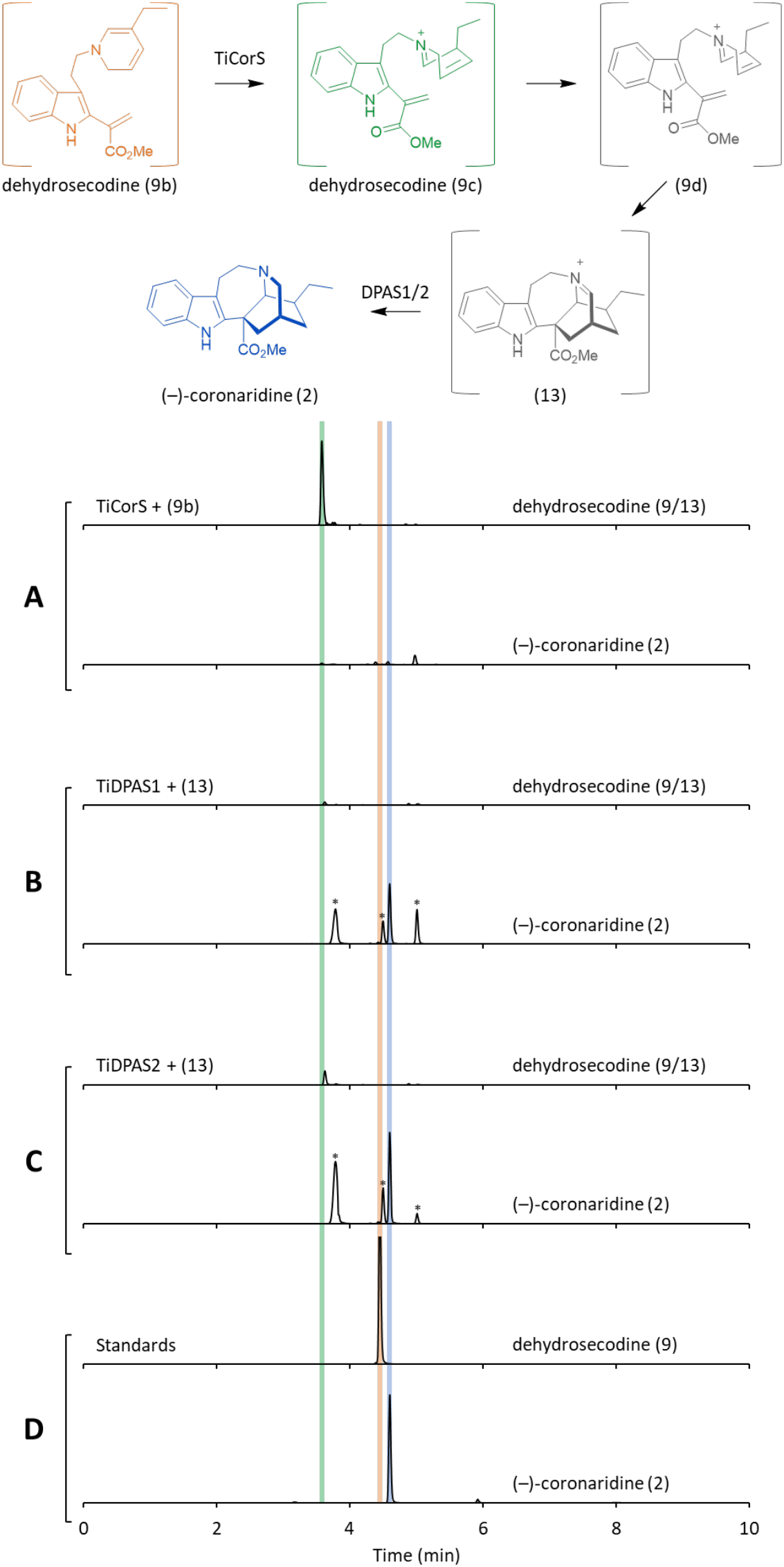
Intermediates 9 and 13 in (−)-coronaridine biosynthesis. **A.** UPLC/MS chromatograms illustrating the formation of **13**(green) from dehydrosecodine isomer **9** (orange) in assays with TiCorS. After 20-minute incubation with TiCorS with **9** to produce **13** (green), either TiDPAS1 (**B**) or TiDPAS2 (**C**) was added to partially purified **13** and NADPH to yield coronaridine (**2**, blue). Authentic standards are shown in (**D**). Peaks marked with * were decomposed during isolation attempts and could not be characterized.

**Fig. S16.**
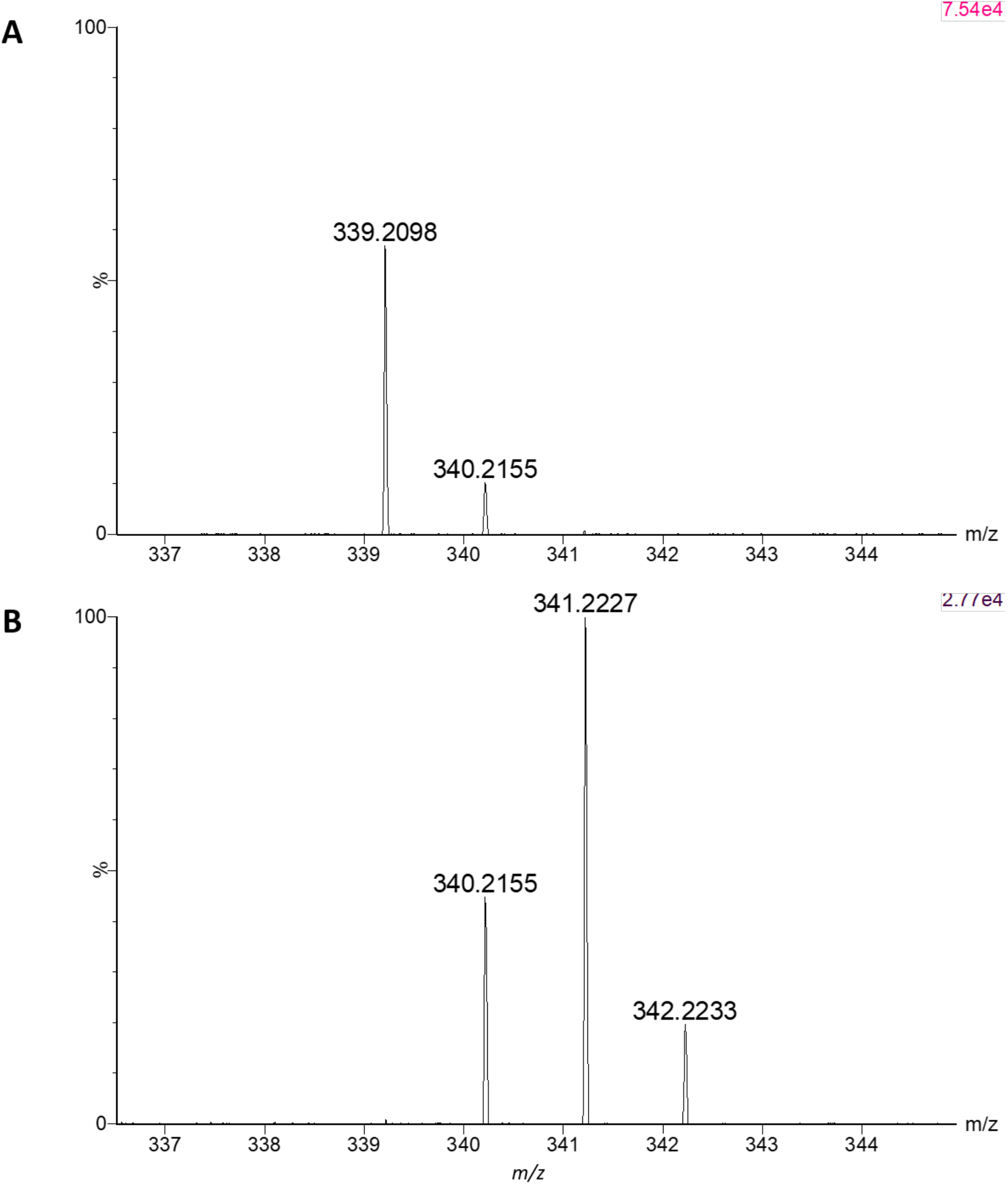
Isotopic labeling of (−)-coronaridine by reaction in D_2_O. **A.** High resolution MS spectra of enzymatically produced coronaridine (**2**) produced in H_2_O. **B.** High resolution MS spectra of enzymatically produced coronaridine (**2**) produced in D_2_O with 1 (*m/z* 340.2) and 2 (*m/z* 341.2) deuterium atoms incorporated.

**Fig. S17.**
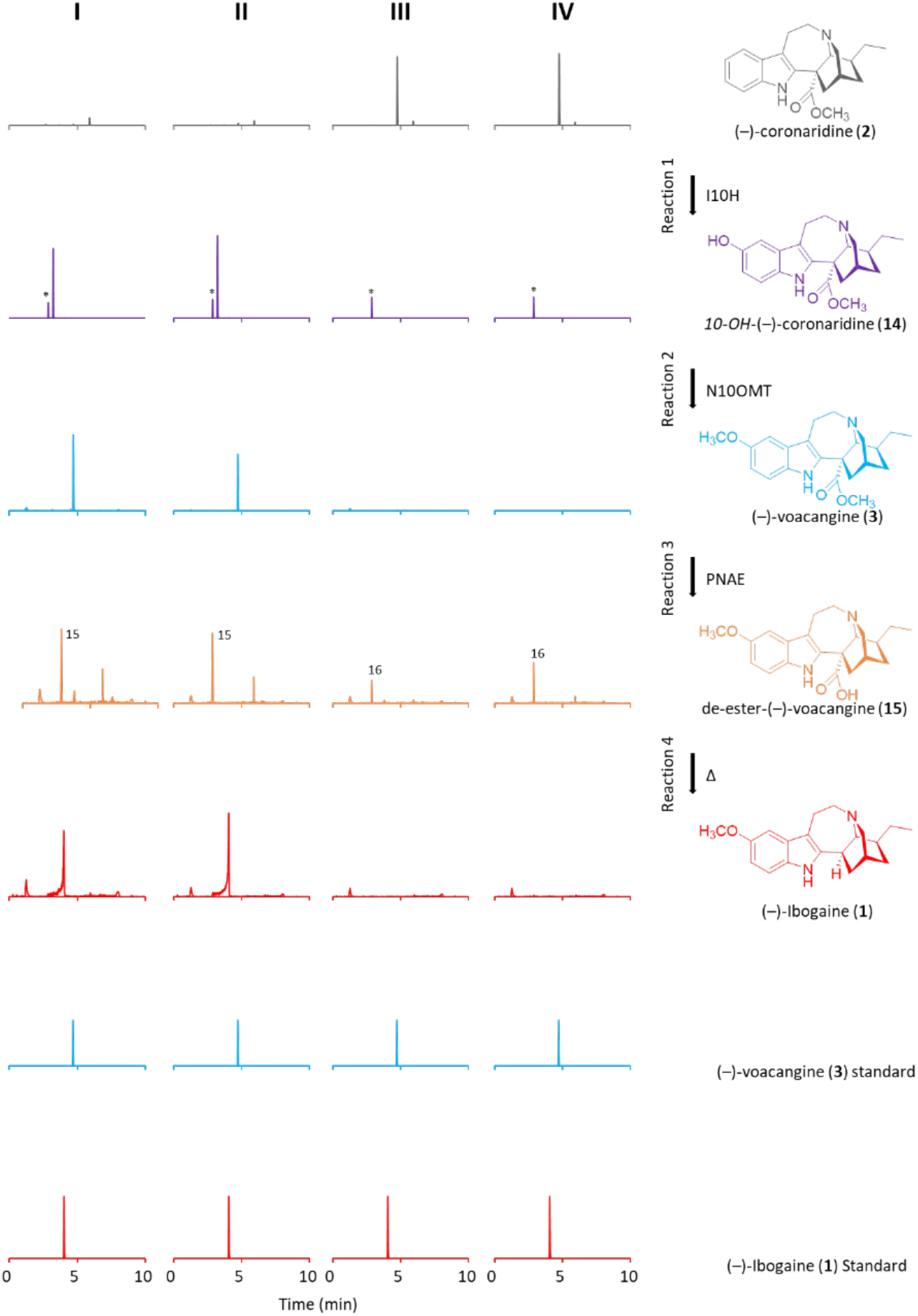
Biocatalytic production of ibogaine. Enzymatically produced coronaridine (**2**) (column I) and authentic (−)-coronaridine (**2**) (column II) incubated with I10H (reaction 1), yields a compound with a mass consistent with (−)-*10*-*OH*-coronaridine (**14**). Addition of N10OMT (reaction 2) to these reactions yields (−)-voacangine (**3**). Subsequent de-esterification of (−)-voacangine (**3**) by TiPNAE (reaction 3) yields (−)-de-ester-voacangine (**15**), and after heating, (−)-ibogaine (**1**). No products were formed when enzymatically produced coronaridine (**2**) (column III) or authentic (−)-coronaridine (**2**) (IV) were incubated with empty vector controls. Peaks marked with an * were products of endogenous yeast enzymes in feeding assays with I10H. De-ester-(−)-coronaridine (**16**) produced via TiPNAE1 in control reactions. These co-elute with de-ester-(−)-voacangine (**15**) and are detected due to MRM inter-channel cross-talk.

**Fig. S18.**
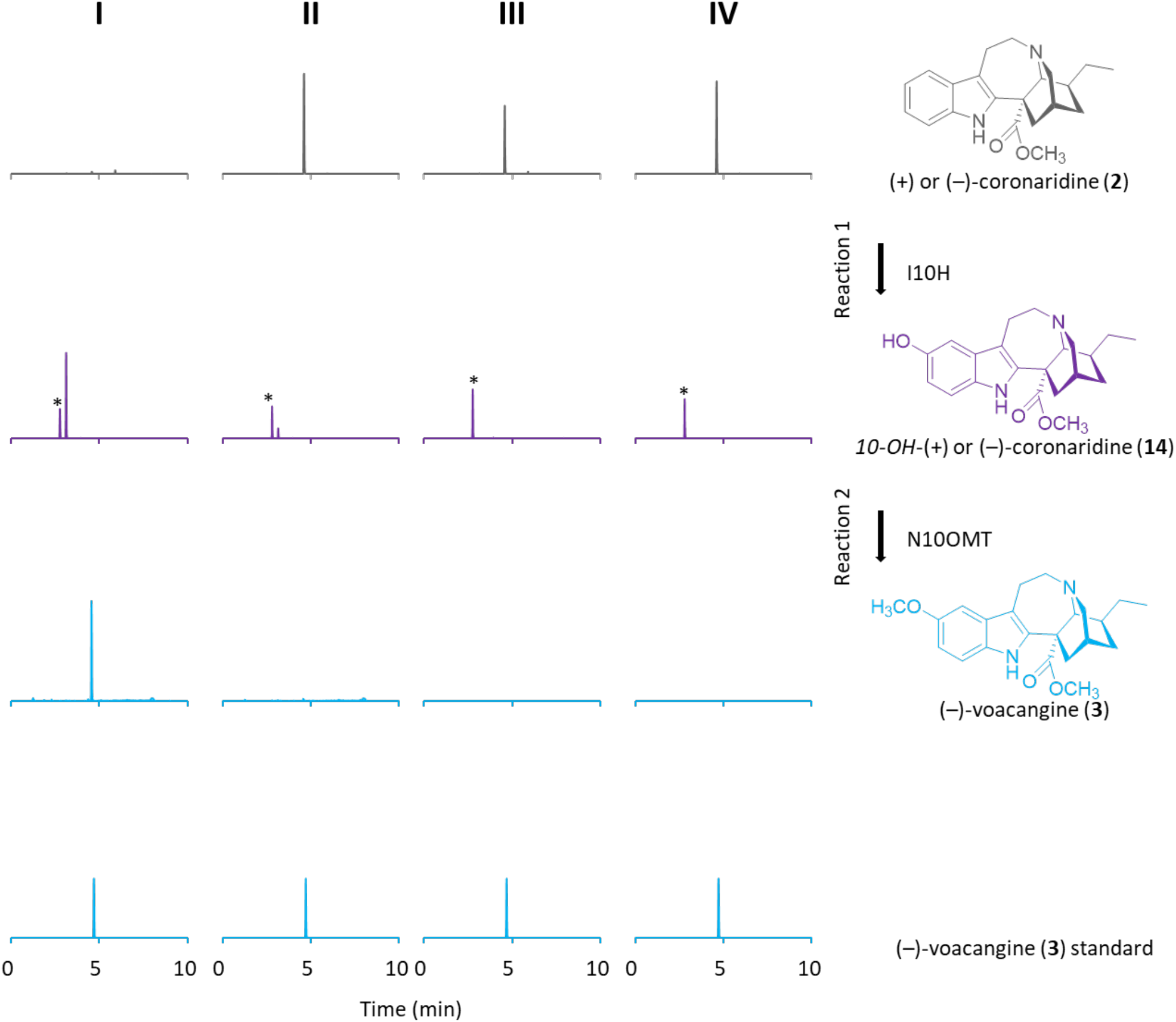
Specificity of I10H and N10OMT for (−)-coronaridine. Enzymatically produced coronaridine (**2**) (column I) incubated with I10H (reaction 1), yields a compound with a mass consistent with 10-*OH*-(−)-coronaridine (**14**) (purple). (+)-Coronaridine (column II) incubated with I10H (reaction 1), yields only trace quantities of 10-*OH*-(+)-coronaridine (**14**) (purple). Addition of N10OMT (reaction 2) to I and II yields (−)-voacangine (**3**) only in I (light blue), showing the specificity of N10OMT for the (−)-coronaridine (**2**) optical series. No products were formed when enzymatically prepared coronaridine (column III) or (+)-coronaridine (**5**) (IV) were incubated with empty vector controls. Peaks marked with * were products of endogenous yeast enzymes present in feeding assays with I10H.

**Fig. S19.**
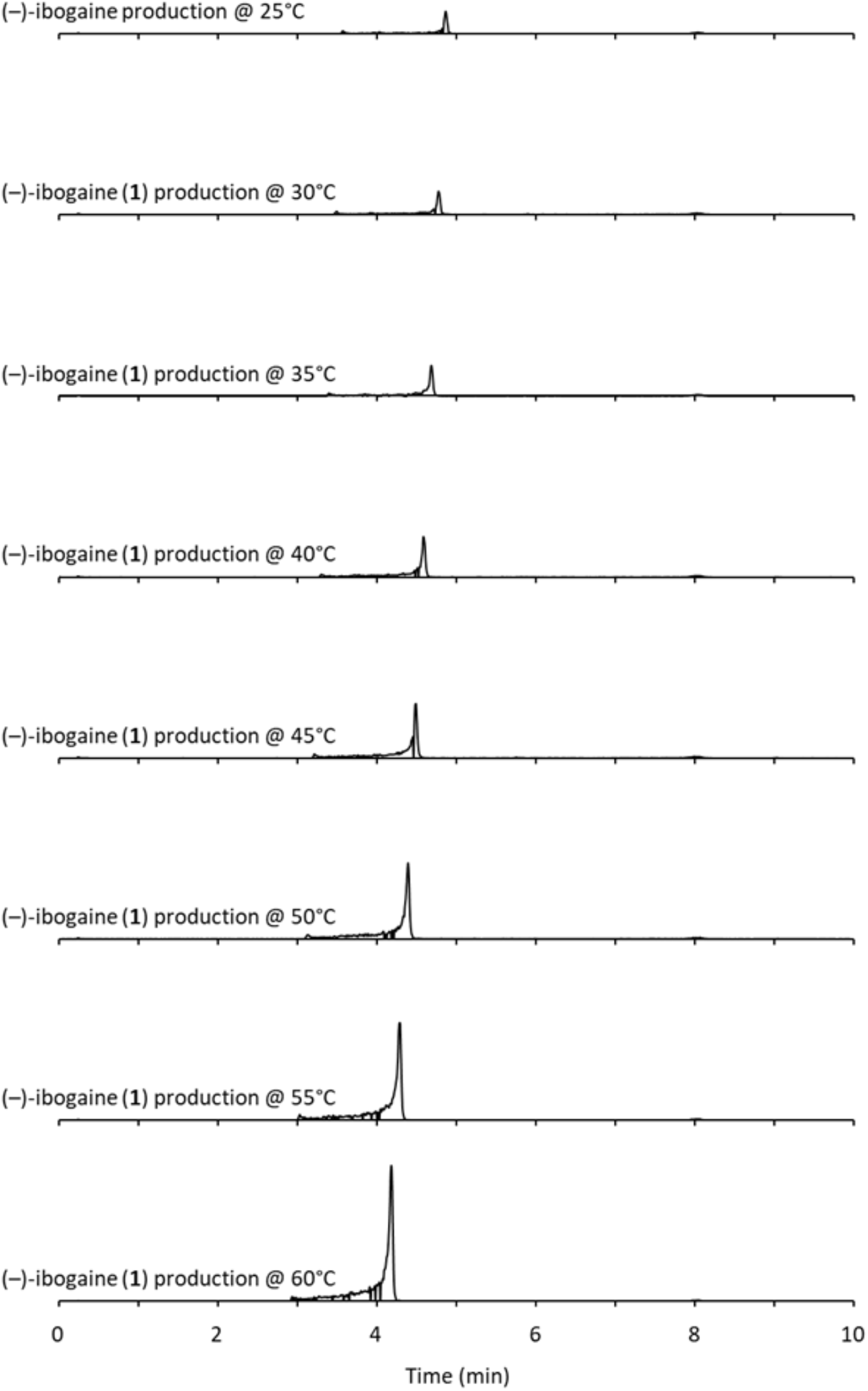
Decarboxylation of (−)-voacangine. After incubation of (−)-voacangine (**3**) with TiPNAE1, the resulting de-esterified (−)-voacangine (**15**) was subjected to increasing temperature on a UPLC column. The decarboxylated product (−)-ibogaine (**1**) increases with increasing temperature.

**Fig. S20.**
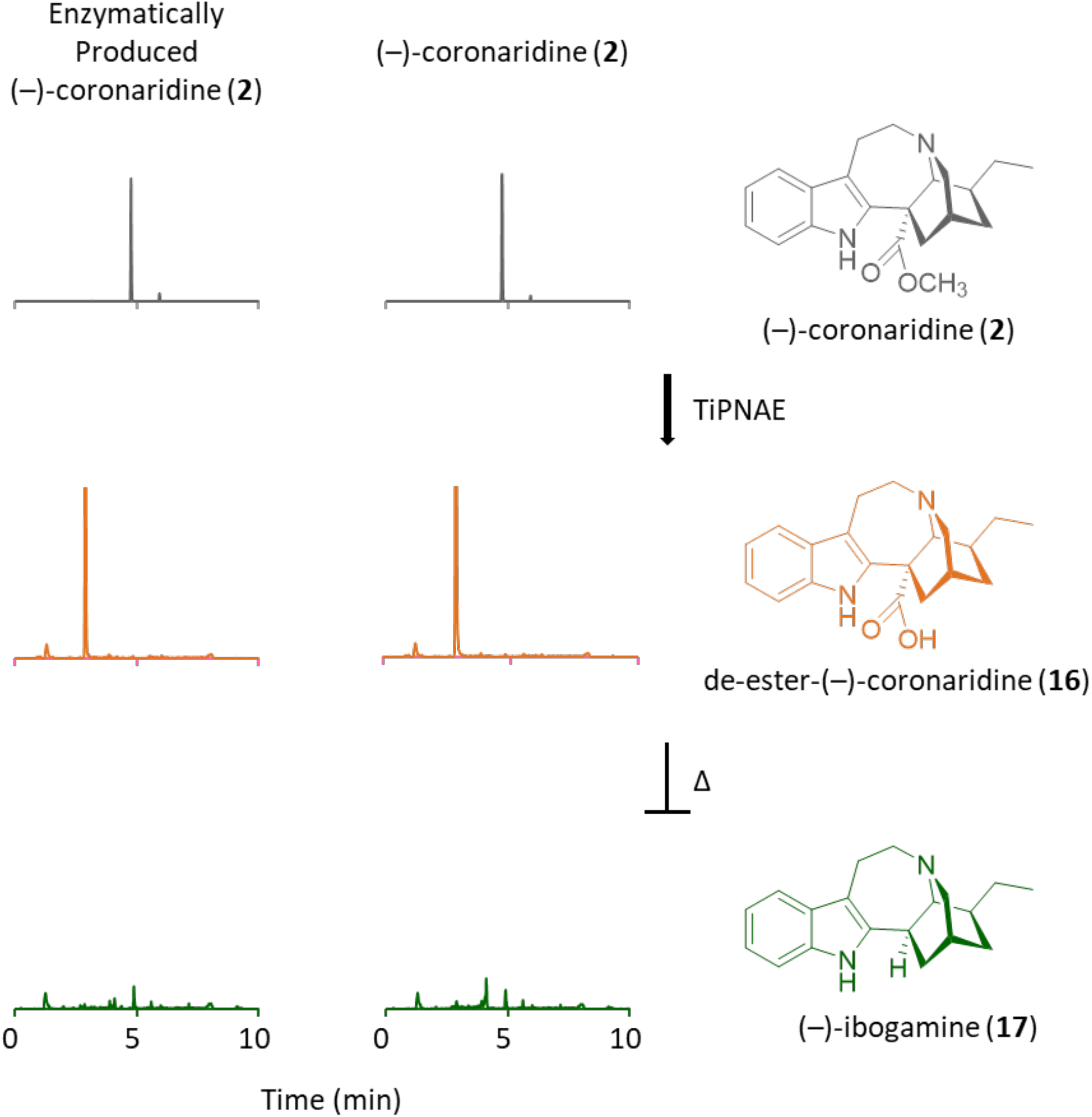
Decarboxylation of coronaridine. Enzymatically produced coronaridine (**2**) and authentic (−)-coronaridine (**2**) are de-esterified by TiPNAE to yield a compound with a mass consistent with de-ester-(−)-coronaridine (**16**); however, heating this product even to 60 °C does not decarboxylate to yield (−)-ibogamine (**17**

**Data S1. (separate file)**

Proteomic analysis of TiPAS1-3 expressed in *N. benthamiana*.

